# *Pten* regulates endocytic trafficking of cell adhesion and signaling molecules to pattern the retina

**DOI:** 10.1101/2022.08.31.506085

**Authors:** Yacine Touahri, Joseph Hanna, Nobuhiko Tachibana, Luke Ajay David, Thomas Olender, Satoshi Okawa, Vorapin Chinchalongporn, Anjali Balakrishnan, Robert Cantrup, Rajiv Dixit, Pierre Mattar, Fermisk Saleh, Yaroslav Ilnytskyy, Monzur Murshed, Paul E. Mains, Igor Kovalchuk, Julie L Lefebvre, Michel Cayouette, Antonio del Sol, Marjorie Brand, Benjamin E Reese, Carol Schuurmans

**Author notes:** Author for correspondence: Carol Schuurmans.

## Abstract

The retina is an exquisitely patterned tissue, with neuronal somata positioned at regular intervals to completely sample the visual field. Cholinergic amacrine cells are spectacular exemplars of precision, distributing in two radial layers and tangentially, forming regular mosaics. Here, we investigated how the intracellular phosphatase Pten and the cell adhesion molecule Dscam cooperate to regulate amacrine cell patterning. Using double mutants to test epistasis, we found that *Pten* and *Dscam* function in parallel pathways to regulate amacrine cell positioning. Mechanistically, *Pten* regulates endocytic remodeling of cell adhesion molecules (Dscam, Megf10, Fat3), which are aberrantly redistributed in *Pten* conditional-knock-out (cKO) amacrine cells. Furthermore, extracellular vesicles derived from multivesicular endosomes have altered proteomes in *Pten*^cKO^ retinas. Consequently, Wnt signaling is elevated in *Pten*^cKO^ retinal amacrine cells, the pharmacological disruption of which phenocopies *Pten*^cKO^ patterning defects. *Pten* thus controls endocytic trafficking of critical cell adhesion/signaling molecules to control amacrine cell spacing.

**HIGHLIGHTS:**

- *Pten* and *Dscam* act in parallel pathways to regulate amacrine cell spacing
- Endocytic remodeling of cell adhesion molecules is perturbed in *Pten*^cKO^ retinas
- Extracellular vesicle content is altered in *Pten*^cKO^ retinas
- Perturbation of Wnt signaling phenocopies defects in amacrine cell positioning

**eTOC BLURB:** Patterns in nature range from stereotyped distributions of colored patches on butterfly wings to precise neuronal spacing in the nervous system. Waddington proposed that built-in constraints canalize developmental patterns. Touahri *et al*. identified *Pten*-mediated endocytic trafficking of cell adhesion/signaling molecules as a novel constraint measure controlling retinal amacrine cell patterning.

## INTRODUCTION

Retinal neurons are spatially organized in two orthogonal dimensions (Wassle, 2004). In the radial axis, neurons form outer nuclear (ONL), inner nuclear (INL), and ganglion cell (GCL) layers. In the tangential dimension, individual types of retinal neurons further organize into regularly spaced arrays known as mosaics (Galli-Resta et al., 2008). This spatial organization allows for the vertical transmission of individual image points and serial light processing and for the parallel processing of visual information to create a composite map of the visual field (Galli-Resta, 2002).

Amacrine cells are born at the apical retinal surface and migrate radially to settle in the inner INL or become displaced in the GCL. Known molecular regulators of radial positioning include the high mobility group-box transcription factor Sox2, which controls the ratio of cholinergic (ChAT^+^) amacrine cells in the GCL versus INL (Whitney et al., 2014). In turn, the atypical cadherin Fat3 controls the bipolar to unipolar transition of radially migrating GABAergic amacrine cell subpopulations (Deans et al., 2011). Mosaic establishment and refinements in the tangential plane occur soon after radial layers form, continuing for the first few postnatal days (Galli-Resta, 2000; Galli-Resta et al., 1997; Reese, 2008; Reese and Galli-Resta, 2002). Two well-studied amacrine cell mosaics are formed by ChAT^+^ and dopaminergic (TH^+^) amacrine cells, which regulate directional sensitivity (Taylor and Smith, 2012) and light adaptation, including transitions between scotopic and photopic vision (Masland, 2012), respectively. ChAT^+^ amacrine cells form an archetypical mosaic, exhibiting exclusion zones that keep them apart from one another (Reese and Galli-Resta, 2002; Rockhill et al., 2000). Such exclusion zones, in the presence of sufficient densities of homotypic cells, ensures a regular patterning within the mosaic (Whitney et al., 2008).

Amacrine cells use several mechanisms to establish mosaics, including selective cell death (if too close to ‘like’ cells), tangential dispersion (moving away from ‘like’ cells), and selective cell fate specification (altering postmitotic phenotypes if too close to ‘like’ cells) (Galli-Resta et al., 2008). Remarkably, each amacrine cell subtype establishes its mosaic pattern independently (Rockhill et al., 2000). ChAT^+^ amacrine cells disperse tangentially (Galli-Resta et al., 1997; Reese et al., 1999; Reese and Tan, 1998) while TH^+^ amacrine cells use lateral inhibition to block the same fate in amacrine cells spaced too close together during early fate determination (Cameron and Carney, 2004; Tyler et al., 2005), followed by selective cell death later on, after cells arrive in their strata (Raven et al., 2003). Interestingly, when a large subset of horizontal cells are depleted at the time of mosaic formation, these cells space themselves further apart with larger exclusion zones, indicating that there are adaptive mechanisms in place to allow complete tiling of the retinal field (Poché et al., 2008).

Strikingly, a computational model based on the generation of diffusible fate-determining factors that negatively feedback to create exclusion zones could accurately simulate mosaic formation (Stenkamp and Cameron, 2002; Tyler et al., 2005). It is therefore surprising that several cell adhesion molecules (CAMs), which traditionally act at the cell surface, control mosaic patterning. Indeed, CAMs have a general role in coordinating neuronal positioning and dendritic arborization patterns within neural circuits (Pollerberg et al., 2013). Not only is the type of CAM expressed important, but also its precise localization, with CAM content tightly regulated by endocytic pathways that traffic new CAMs to the cell membrane, and remove membrane CAMs, either to be degraded or transferred to extracellular vesicles (EVs) for secretion. For instance, in *Drosophila*, Neuroglian, an L1-type CAM, is redistributed from the plasma membrane to endosomes during dendritic pruning in a subset of dendritic arborization (da) neurons (Zhang et al., 2014). Similarly, Downs syndrome cell adhesion molecule (Dscam), which has over 38,000 isoforms in the fly, covers da neuronal dendrites, but this coating is reduced during dendritic tiling (Meltzer and Schuldiner, 2022). In contrast, in mammals, Dscam maintains TH^+^ amacrine cell mosaics by acting as a “non-stick coating”, blocking adhesive signals (Fuerst et al., 2012; Garrett et al., 2018; Keeley et al., 2012). Thus, in the absence of *Dscam*, instead of TH^+^ amacrine cell dendrites forming continuous nets that uniformly cover the retinal surface, dendrites hyperfasciculate, which secondarily degrades the somal mosaic. Other CAMs involved in amacrine cell spacing include *Megf10*, a transmembrane molecule, which mediates homotypic repulsion between ChAT^+^ amacrine cells to establish somal mosaics (Kay et al., 2012). In addition, differential expression of γ-protocadherin (*Pcdhg*) isoforms confers ChAT^+^ amacrine cell self-recognition such that dendrites on the same amacrine cell do not ectopically adhere together (Lefebvre et al., 2012).

Strikingly, *Phosphatase and tensin homolog* (*Pten*) also regulates radial and tangential positioning of amacrine cells and prevents the fasciculation of TH^+^ neurites (Cantrup et al., 2012). How an intracellular phosphatase mediates cell recognition between cells spaced a large distance apart, and coordinates non-overlapping dendrite extensions, is unclear. Here, we reveal that *Pten* controls amacrine cell and process positioning by regulating two modes of endosomal trafficking: (1) the recycling of CAMs to their proper location on the cell membrane, and (2) the coordinated exocytosis of important signaling molecules, including Wnt pathway molecules, which normally traffic from multivesicular endosomes into secreted EVs. Finally, using a pharmacological inhibitor of Wnt signaling, we partially phenocopy ChAT^+^ amacrine cell spacing defects observed in *Pten* cKOs. Taken together, *Pten-*regulated endosomal trafficking functions as a built-in constraint mechanism to ensure that cell adhesion and signaling molecules are properly distributed in, and secreted by, amacrine cells, to ultimately coordinate retinal patterning.

## RESULTS

### *Pten* and *Dscam* are co-expressed in dopaminergic and cholinergic amacrine cells

Both *Pten* (Cantrup et al., 2012) and *Dscam* (Fuerst et al., 2008) are required for the proper spatial distribution of TH^+^ amacrine cells, with the loss of these two genes leading to TH^+^ amacrine cell clumping and neurite fasciculation. To determine whether and how these two genes might cooperate, we first analyzed their expression in the retina using RNAscope to localize their transcripts. In postnatal day (P) 7 wild-type retinas, at a developmental stage when amacrine cell differentiation is complete and mosaic refinement is in progress, *Pten* and *Dscam* transcripts were both detected in the INL and GCL in the retina (Figure 1A). However, while *Dscam* mRNA puncta were restricted to the inner half of the INL, where amacrine cells reside, *Pten* transcripts were more widely distributed and also detected in the upper INL, which contains the somata of bipolar, horizontal, and Müller glial cells (Figure 1A). Higher magnification images confirmed the overlap in *Pten* and *Dscam* transcript localization in the same cell, indicating that these two genes are indeed co-expressed in inner INL cells at P7 (Figure 1A).

**Figure 1.**
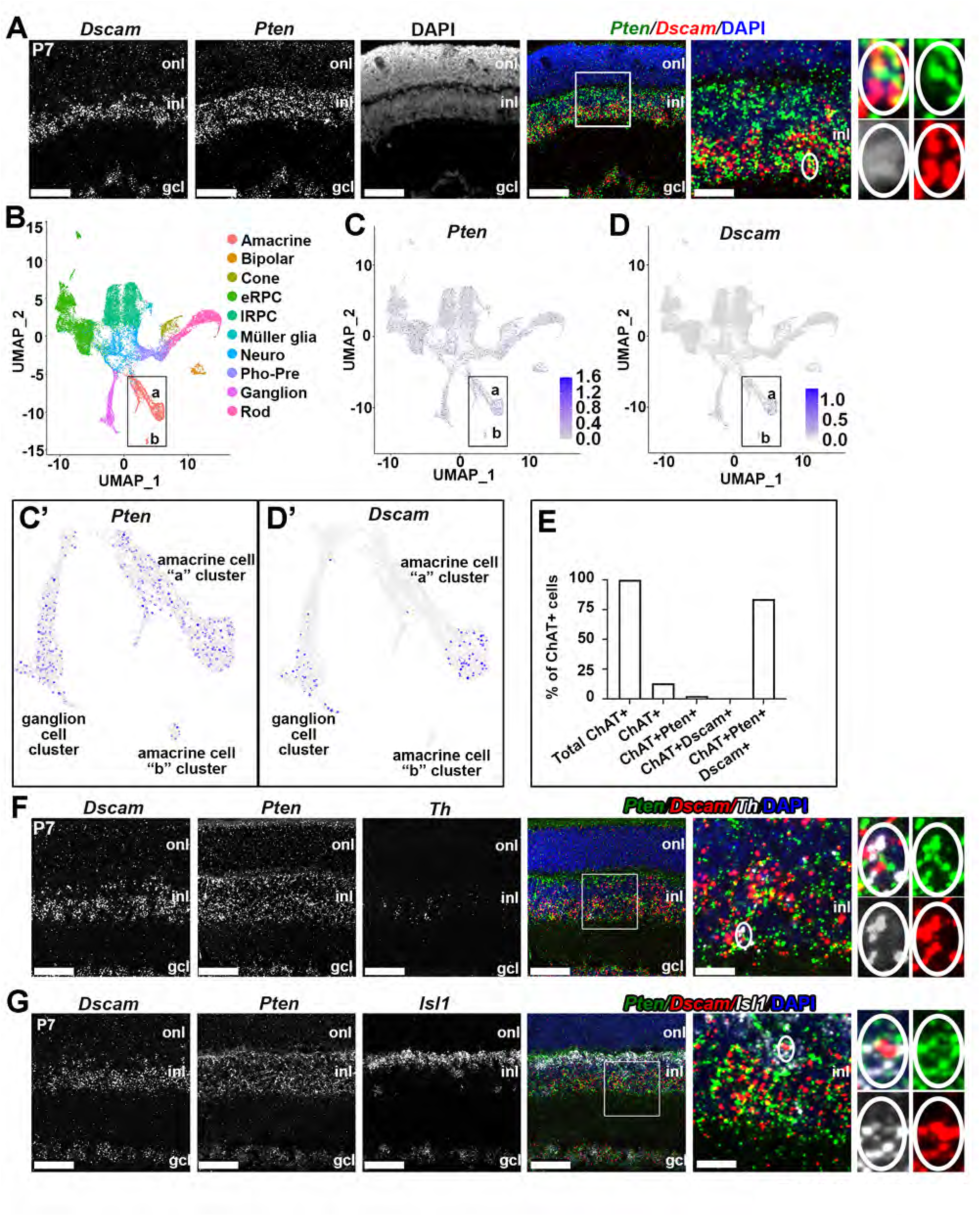
*Pten* and *Dscam* are co-expressed in dopaminergic and cholinergic amacrine cells. (A) RNAscope *in situ* hybridization of P7 retinas, showing co-localization of *Dscam* and *Pten* transcripts in the lower INL. High magnification image of a single cell to the right show transcripts for *Dscam* and *Pten* are located in the same cell. (B-E) UMAP plot of scRNA-seq data from E11-P14 retinal cells, with cluster identities assigned using signature gene expression, and indicated with a color code (B) (data mined from (Clark et al., 2019)). Expression of *Pten* (C,C’) and *Dscam* (D,D’) in retinal cell clusters, with high magnification images of the amacrine cell “a” and “b” clusters in C’ and D’. Quantification of the percentage of ChAT^+^ cells positive for *Pten*, positive for *Dscam* and positive for both *Pten and Dscam* in the scRNAseq dataset (Clark et al., 2019) (E). (F,G) RNAscope in situ hybridization of P7 retinas, showing co-labeling of *Dscam, Pten* and *Th* (F) and *Dscam, Pten* and *Isl1* (G). High magnification images of single cells to the right show transcripts for *Dscam, Pten* and either *Th* or *Isl1* are located in the same cell. Blue is DAPI counterstain. gcl, ganglion cell layer; inl, inner nuclear layer; onl, outer nuclear layer. Scale Bar: 50μm in first four panels, and 10μm in high magnification images in the fifth panel over.

To provide further support for the co-expression of *Pten* and *Dscam* in amacrine cells at single-cell resolution, we mined available scRNAseq data pooled from 10 stages of retinal development between embryonic day (E) 11, at the start of neurogenesis, and P14, when retinal neurogenesis is complete (Clark et al., 2019). High dimensional uniform manifold approximation and projection (UMAP) plots were generated, and signature genes from Clark et al. (2019) were used for cluster annotation, identifying 10 unique cellular clusters that included early (eRPC) and late (lRPC) retinal progenitor cells and eight mature retinal cell types (amacrine, bipolar, cone, Müller glia, photoreceptor precursors, neurogenic precursors (‘neuro’), ganglion and rod cells; Figure 1B). The amacrine cell cluster included one larger (“a”) and one smaller (“b”) cluster, both of which expressed *Pten* (Figure 1C,C’). In contrast, *Dscam* expression was largely restricted to ganglion cells and amacrine cells in “cluster a” (Figure 1D,D’), at least at this sequencing depth.

We next quantified the percentage of ChAT^+^ and TH^+^ amacrine cells that expressed only *Pten*, only *Dscam*, or both *Pten* and *Dscam*. Of the 119/16560 sequenced retinal cells from the Clark et al. (2019) study that expressed ChAT, 84% co-expressed both *Pten* and *Dscam* (Figure 1E). Surprisingly, only 35 TH^+^ amacrine cells were sequenced in this data set, and of these, none was positive for *Pten* and *Dscam*. However, the lack of double or triple positive TH^+^ amacrine cells could reflect sequencing depth and the lack of maturity of most cells in the pool sequenced between E11-P14. For that reason, we also used RNAscope to examine the co-expression of *Pten* and *Dscam* with *Th* or *Isl1*, a transcription factor that is expressed in bipolar cells in the outer INL and ChAT^+^ amacrine cells in the inner INL (Elshatory et al., 2007). In P7 retinas, *Th* transcripts were sparsely distributed in the INL (Figure 1F), as expected given that TH^+^ amacrine cells are one of the most sparsely distributed cell types present, amounting to less than one-hundredth of one percent of all retinal neurons (Whitney et al., 2009). However, we were able to localize a few triple-labeled TH^+^ amacrine cells that co-expressed *Pten*, *Dscam,* and *Th* (Figure 1F). In contrast, *Isl1* transcripts were abundant in the upper INL, presumably in bipolar cells, and in the inner half of the INL, where *Dscam* and *Pten* transcripts are concentrated (Figure 1G). ChAT^+^ amacrine cells that co-expressed *Pten, Dscam,* and *Isl1* were also detected (Figure 1G). Thus, *Pten* and *Dscam* are co-expressed in a few TH^+^ amacrine cells, and in the majority of early ChAT^+^ amacrine cells.

### Generation of *Pten* and *Dscam* single and double mutant embryos reveals postnatal lethality

To determine whether *Pten* and *Dscam* function in the same or in parallel pathways to control amacrine cell positioning, we performed double mutant analysis, comparing the phenotypes of double mutants to single mutants. To study *Dscam* mutants, we used homozygotes carrying a del17 null allele with a 38 bp deletion in exon 17 (hereafter *Dscam*^del17^), which causes a frameshift mutation and generates a premature stop codon (Fuerst et al., 2008). For *Pten*, we generated a conditional knock-out (cKO) by crossing mice carrying a *Pten*^fl/fl^ allele with floxed exons 4 and 5 (Backman et al., 2001), with a *Pax6* α-enhancer/P0 promoter::*Cre-IRES-GFP* driver line (Marquardt et al., 2001). *Pax6::Cre* promotes recombination beginning at E10.5 in RPCs of the peripheral retina, and hence all progeny neurons, including amacrine cells are mutated for *Pten* in retinal domains where Cre is expressed (Marquardt et al., 2001).

We crossed *Pten^fl/+^;Pax6::Cre^+^;Dscam^del17/+^*males with *Pten^fl/+^;Dscam^del17/+^* females to generate ‘wild-type’ (all non-mutant genotypes), *Pten*^cKO^ and *Dscam*^del17^ single mutants and *Pten*^cKO^*;Dscam*^del17^ double mutants. We collected 56 litters for a total of 376 live P14 pups and compared the acquired to expected genotypes (Figure S1A). Several genotypes were under-represented based on theoretical Mendelian ratios, including all homozygous *Dscam*^del17^ combinations (Figure S1B), consistent with the prior claim that homozygous *Dscam* null mutants are perinatally lethal (Amano et al., 2009). However, *Dscam*^del17^ lethality was not completely penetrant, allowing us to analyse surviving pups.

### Disruption of the inner nuclear layers is exacerbated in *Pten*^cKO^*;Dscam*^del17^ double mutants

We first examined the distribution of amacrine cells in the radial axis in P14 wild-type, *Pten*^cKO^ or *Dscam*^del17^ single mutants, and *Pten*^cKO^*;Dscam*^del17^ double mutant retinas (Figure 2). In P14 wild-type retinas, DAPI staining clearly delineated the three retinal layers (ONL, INL, GCL; Figure 2A). In contrast, in *Pten*^cKO^ retinas, there was a disruption of retinal layering, most notably affecting the INL and GCL, with DAPI^+^ nuclei located in an expanded IPL, as previously reported (Cantrup et al., 2012; Sakagami et al., 2011), which is normally a cell free-zone in wild-type retinas (Figure 2A). Similarly, in P14 *Dscam*^del17^ retinas, the INL and GCL were less tightly packed, although they remained as distinct layers, and DAPI^+^ nuclei were also detected in the IPL, resembling the *Pten*^cKO^ phenotype (Figure 2A). Finally, in P14 *Pten*^cKO^*;Dscam*^del17^ double mutant retinas, the IPL was even more grossly expanded, and the cell-free zone in the IPL was even less distinct compared to wild-type and single mutants, with cells scattered throughout the INL, IPL, and GCL (Figure 2A).

**Figure 2.**
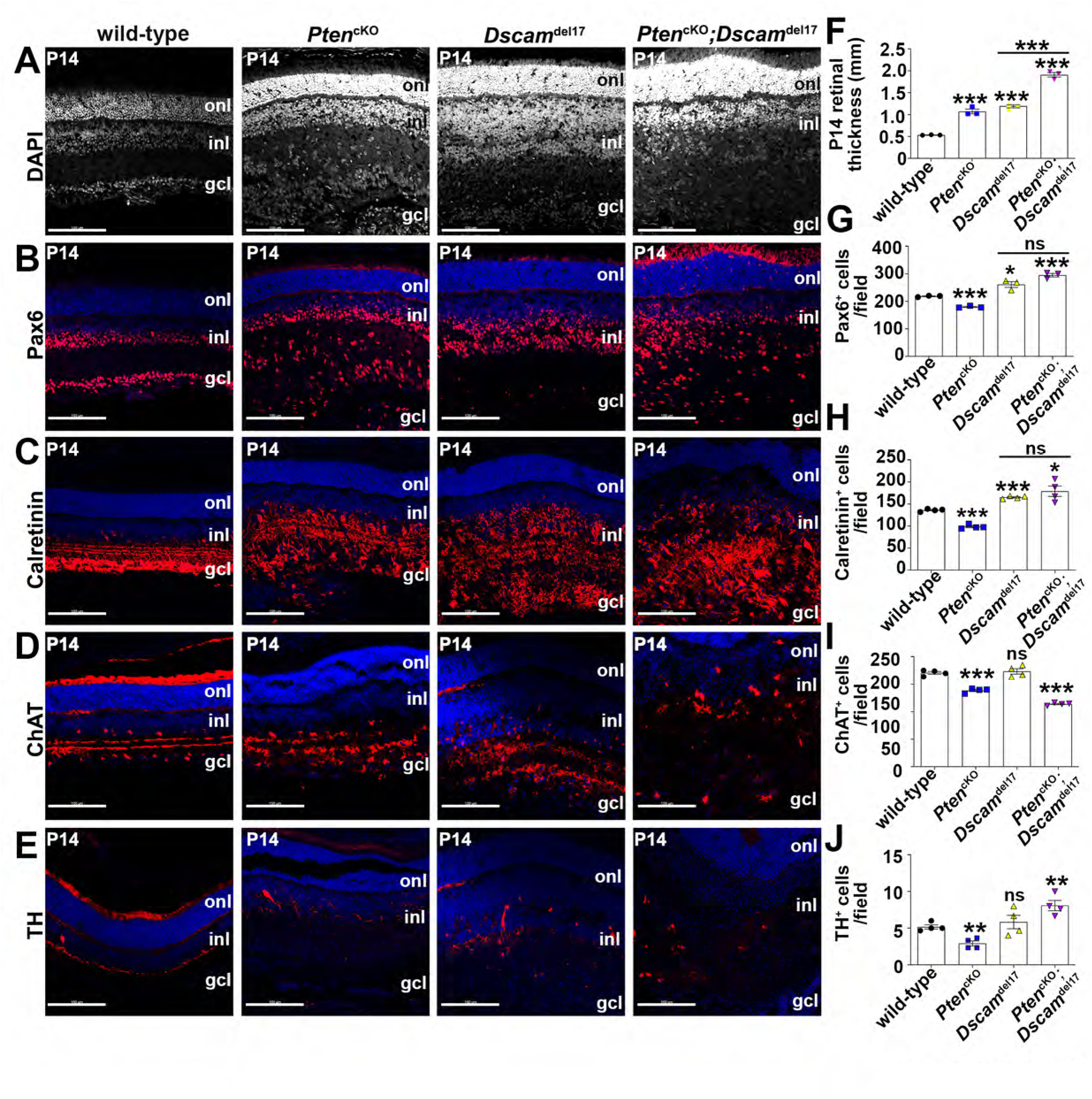
Defects in amacrine cell genesis and radial positioning in *Pten* and *Dscam* single and double mutant retinas. (A) DAPI staining of transverse sections through the retinas of P14 wild-type, *Pten*^cKO^, *Dscam*^del17^, and *Pten*^cKO^*;Dscam*^del17^ retinas. (B-E) Immunolabeling of P14 wild-type, *Pten*^cKO^, *Dscam*^del17^, and *Pten*^cKO^*;Dscam*^del17^ retinas with Pax6 (B), Calretinin (abbreviated CalR, C), and Choline acetyltransferase (abbreviated ChAT, D) and Tyrosine hydroxylase (abbreviated Th, E). (F-J) Quantification of retinal thickness (F), Pax6^+^ (G), Calretinin^+^ (H), ChAT^+^ (I) and TH^+^ (J) cells per field of view. gcl, ganglion cell layer; inl, inner nuclear layer; onl, outer nuclear layer. p values are denoted as follow: <0.05 *, <0.01 **, <0.001 ***. Scale Bar: 200μm. See also Figure S1, S2.

To quantify these disturbances, we measured retinal thickness, revealing 2-fold (n=3; p<0.001) and 2.2-fold (n=3; p<0.001) increases in P14 *Pten*^cKO^ and *Dscam*^del17^ retinas, respectively (Figure 2F). More strikingly, in *Pten*^cKO^*;Dscam*^del17^ retinas, there was an additive effect with a 3.6-fold increase in retinal thickness relative to wild-type retinas (n=3; p=0.0001). Comparisons of the double mutant to *Pten*^cKO^ and *Dscam*^del17^ single mutant retinas also revealed 1.6-fold (n=3; p<0.001) and 1.8-fold (n=3; p<0.001) increases, respectively (Figure 2F). The severity of the double mutant phenotype developed over time, since at P7, there were 1.2-fold (n=3; p<0.0001) and 1.4-fold (n=3; p<0.005) expansions in *Pten*^cKO^ and *Dscam*^del17^ single mutant retinas, respectively (Figure S2A,B). There was a similar 1.4-fold increase in the thickness of P7 *Pten*^cKO^*;Dscam*^del17^ double mutant retinas, with some cells scattered throughout the INL, IPL and GCL, but this expansion was comparable to *Dscam*^del17^ retinas (Figure S2D,E). Thus, exacerbation of retinal layer expansion in *Pten*^cKO^*;Dscam*^del17^ retinas occurs within the time frame of P7 to P14.

Taken together, these data suggest that *Pten* and *Dscam* act in an additive fashion, implying parallel pathways, to control retinal thickness, and indicate that they are partially redundant for the establishment of pattern formation in the radial axis of the retina.

### *Pten* acts upstream as a negative regulator of *Dscam* in the control of amacrine cell number and positioning

We previously demonstrated that not only was there a disruption of TH^+^ amacrine cell positioning in *Pten*^cKO^ retinas, but also a reduction in amacrine cell number (Tachibana et al., 2016). Conversely, more AII amacrine cells are present in *Dscam*^del17^ retinas (Fuerst et al., 2009), although pan-amacrine cell markers were not examined. Here we compared amacrine cell numbers in P14 wild-type, *Pten*^cKO^ and *Dscam*^del17^ single mutants and *Pten*^cKO^*;Dscam*^del17^ double mutant retinas, quantifying the number of cells expressing Pax6, a pan-amacrine cell marker (Haverkamp and Wassle, 2000; Yan et al., 2020), as well as calretinin (Calb2), which labels several GABAergic amacrine cells, including the A17 amacrine cell, one of over 60 amacrine cell subtypes recently identified (Haverkamp and Wassle, 2000; Yan et al., 2020). In P14 *Pten*^cKO^ retinas, there was a 1.2-fold reduction in the number of Pax6^+^ amacrine cells compared to wild-type (n=3; p=0.0001; Figure 2B,G), consistent with previous reports (Tachibana et al., 2016). In contrast, 1.2-fold (n=3; p<0.05) and 1.3-fold (n=3; p<0.001) more Pax6^+^ amacrine cells were detected in P14 *Dscam*^del17^ single and *Pten*^cKO^*;Dscam*^del17^ double mutant retinas, respectively, compared to wild-type (Figure 2B,G). Notably, Pax6^+^ amacrine cell number was not statistically different when comparing P14 *Dscam*^del17^ and *Pten*^cKO^*;Dscam*^del17^ retinas (n=3; p>0.05; Figure 2G). A similar pattern was observed when examining calretinin^+^ amacrine cells, with a 1.4-fold reduction in calretinin^+^ amacrine cells in *Pten*^cKO^ retinas compared to wild-type retinas (n=3; p<0.0001), versus a 1.2-fold increase in *Dscam*^del17^ retinas (n=3; p<0.0001) and a 1.3-fold increase in *Pten*^cKO^*;Dscam*^del17^ retinas (n=3; p<0.05; Figure 2C,H). Thus, the increase in amacrine cell number in *Dscam*^del17^ mutants is epistatic to the decrease seen in *Pten*^cKO^ retinas. With respect to amacrine cell number, *Pten* and *Dscam* are therefore in a different relationship than for layer thickness, with *Pten* acting upstream of *Dscam* as a negative regulator. These data suggest that *Pten* loss perturbs Dscam expression (as verified below). The *Pten*^cKO^ amacrine cell number phenotype is thus dependent, at least in part, on the presence of wild-type Dscam.

We next assessed the stratification of amacrine cell processes in the radial plane. Retinas were thicker in all *Pten* and *Dscam* single and double mutants, an expansion that was primarily observed in the IPL. Within the IPL, amacrine cells elaborate dendritic arbors that stratify in unique radial patterns, forming five discrete sublaminae (s1-s5) (Famiglietti and Kolb, 1976; Wassle, 2004). Calretinin^+^ amacrine cells normally arborize in three strata in the IPL (s1-s3), as seen in P14 wild-type retinas (Figure 2C). The patterning of calretinin^+^ amacrine cell IPL dendrites was disrupted in P14 *Pten*^cKO^ and *Dscam*^del17^ single mutant retinas, with only a shadow of the three layers detected, and scattered cells detected in this synaptic zone (Figure 2C). In contrast, in P14 *Pten*^cKO^*;Dscam*^del17^ retinas, there was no stratification visible in the IPL, and calretinin^+^ amacrine cells were scattered throughout the IPL (Figure 2C).

Similarly, the expression of ChAT, a marker for cholinergic starburst amacrine cells, which arborize in s2 and s4 in the IPL, was also disrupted in P14 *Pten*^cKO^ and *Dscam*^del17^ retinas, with indistinct s2 and s4 strata, while in *Pten*^cKO^*;Dscam*^del17^ retinas, there was no obvious ChAT^+^ stratification visible in the IPL (Figure 2D). Finally, TH^+^ amacrine cell dendrites normally mono-stratify in s1, as observed in P14 wild-type retinas, and while the TH^+^ dendritic IPL pattern was visible in s1 in P14 *Pten*^cKO^ and *Dscam*^del17^ single mutant retinas, TH labeling was discontinuous (Figure 2E). In contrast, no stratification was visible in *Pten*^cKO^*;Dscam*^del17^ retinas (Figure 2E), indicating additivity.

Finally, we analysed ChAT^+^ and TH^+^ cell numbers, revealing that both were reduced in *Pten*^cKO^ and not in *Dscam*^del17^ retinas (Figure 2D,E). Conversely, while ChAT^+^ cell number was reduced in *Pten*^cKO^*;Dscam*^del17^ retinas compared to wild-type and *Pten*^cKO^ (n=3; p<0.05; Figure 2D,I), TH^+^ amacrine cell number was higher compared to wild-type and *Pten*^cKO^ (n=3; p<0.05; Figure 2E,J). In summary, amacrine cell number and radial layering were strikingly perturbed in all three mutants, with defects most evident in IPL patterning, and amacrine cell organization in *Pten*^cKO^*;Dscam*^del17^ double mutant retinas. As the cell number and stratification phenotypes were more striking when both genes were mutated, these data suggest that *Pten* and *Dscam* act in parallel pathways to control amacrine cell differentiation, retinal lamination, and IPL stratification, and indicates that both genes are major players in this process.

### *Pten* and *Dscam* act in parallel pathways to control cholinergic amacrine cell distribution in the tangential plane

We found that *Pten* and *Dscam* are co-expressed in most ChAT^+^ amacrine cells (Figure 1E), and they contribute in parallel pathways to establish the radial organization of these cells and their processes. These findings prompted us to next ask whether these two genes are involved in the spatial organization of ChAT^+^ amacrine cells in the tangential axis. We first examined the tangential spatial organization of ChAT^+^ amacrine cells in *Pten*^cKO^ retinas, as we had previously only assessed TH^+^ cells (Cantrup et al., 2012). Presumptive ChAT^+^ amacrine cells were labelled with Islet1 at P7 (Elshatory et al., 2007), and with ChAT at P14 and P21. We used two spatial statistical measures to assess cellular order; Voronoi domain and nearest neighbor analyses, each derived from the Delauney tessellation of the field (Raven and Reese, 2002), as previously implemented (Cantrup et al., 2012). The frequency of Voronoi domain area sizes (being the territories closer to each cell than to any of their neighbors) and the nearest neighbor distances were plotted, and variability was assessed using a regularity index (RI).

In P7 wild-type retinas, Islet1^+^ amacrine cell somata in the INL were distributed in a regular, patterned array, indicated by the limited variability of Voronoi domain areas (Figure S3A-B), as previously reported (Galli-Resta et al., 1997). In contrast, in P7 *Pten*^cKO^ retinas, the Voronoi domain regularity index (VDRI) showed a 1.4-fold reduction, indicative of less regular spacing (n=3; p<0.001; Figure S3D). A comparable 1.2-fold reduction was also seen in the nearest neighbor regularity index (NNRI) of Islet1^+^ amacrine cells in P7 *Pten*^cKO^ retinas (Figure S3A-H). Similarly, at P14, the VDRI for ChAT^+^ amacrine cells was reduced 1.3-fold in *Pten*^cKO^ compared to wild-type retinas (n=3; p<0.001; Figure 3A,B,E), and at P21 there was a 1.7-fold reduction in *Pten*^cKO^ retinas (n=3; p<0.0001; Figure S3I-J,L). Finally, the NNRI was reduced 1.1-fold in P14 *Pten*^cKO^ retinas compared to wild-type, nearly reaching significance (n=3; p=0.057; Figure 3A,B,F) and 1.5-fold at P21 (n=3; p<0.01; Figure S3Q). ChAT^+^ amacrine cell tangential spatial organization is thus disrupted in *Pten*^cKO^ retinas from as early as P7.

**Figure 3.**
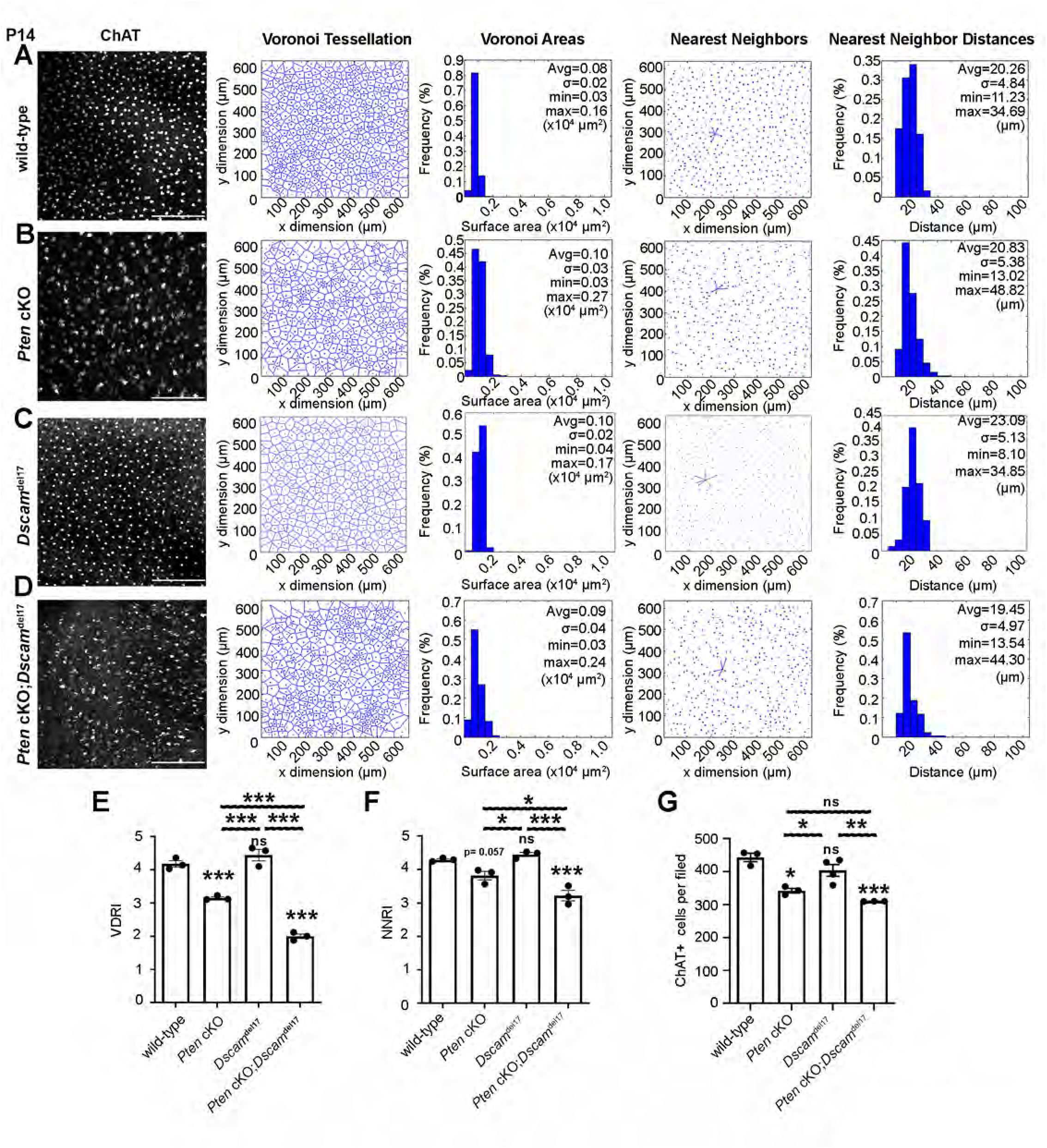
Aberrant cholinergic amacrine cell mosaics in *Pten* and *Dscam* single and double mutant retinas. ChAT immunolabeling and mosaic analysis of P14 wild-type (A), *Pten*^cKO^ (B), *Dscam*^del17^ (C) and *Pten;Dscam* DKO (D) retinal flatmounts showing Voronoi Tessellations, Nearest neighbor plots and frequency distributions of nearest neighbor distances for ChAT^+^ INL amacrine cells. Calculation of Voronoi domain (E) and Nearest Neighbor (F) regularity indices and ChAT^+^ cell number per field (G). p values are denoted as follow: <0.05 *, <0.01 **, <0.001 ***. Scale Bar: 200μm. See also Figure S3.

Our data indicate that the majority of ChAT^+^ amacrine cells co-express *Pten* and *Dscam,* so we compared *Pten*^cKO^ mutants to *Dscam*^del17^ single or *Pten*^cKO^*;Dscam*^del17^ double mutant retinas. A previous study reported that *Dscam*^del17^ retinas have normal cholinergic arrays (Fuerst et al., 2009), and indeed, no significant difference was found between *Dscam*^del17^ and wild-type controls for either the VDRI or NNRI (Figure 3A,C,E,F). In striking contrast, the VDRI for ChAT^+^ amacrine cells was reduced 2.1-fold in P14 *Pten*^cKO^*;Dscam*^del17^ double mutant compared to wild-type retinas (n=3; p<0.001), and 1.6-fold compared to *Pten*^cKO^ retinas (n=3; p<0.001; Figure 3E). Similar to *Pten*^cKO^ retinas, the NNRI for ChAT^+^ amacrine cells in P14 *Pten*^cKO^*;Dscam*^del17^ retinas was reduced 1.3-fold compared to wild-type retinas (n=3; p<0.01), and 1.6-fold compared to *Pten*^cKO^ retinas (n=3; p<0.001; Figure 3F). We confirmed in these retinal wholemounts the differences in cell number estimated from the retinal sections above (Figure 2I), finding that ChAT^+^ cell density was significantly reduced in both *Pten*^cKO^ and *Pten*^cKO^*;Dscam*^del17^ retinas compared to wild-type (Figure 3G).

Collectively, these data suggest that *Pten* and *Dscam* act in parallel in a partially redundant fashion to regulate the spatial organization of ChAT^+^ amacrine cells in the tangential plane However, *Pten* is the major player and *Dscam* a minor player as *Dscam* loss of function only affects ChAT^+^ tangential distribution in the context of *Pten*^cKO^.

### Abnormal expression of cell adhesion molecules in *Pten^cKO^* cholinergic amacrine cells

To better understand how Pten, an intracellular phosphatase, and CAMs, which decorate the plasma membrane, both control amacrine cell spacing, we focused on the endocytic compartment as a potential link. CAM expression on the plasma membrane is achieved by a balance of active trafficking of newly translated CAMs to the cell surface and constant remodeling by endocytosis through recycling endosomes (Kawauchi, 2012). Pten also localizes to endosomal membranes, and shares sequence homology with auxilin, which is essential for endocytosis (Naguib et al., 2015). We therefore questioned whether in the absence of *Pten*, the expression of CAMs, which is tightly regulated by recycling endosomes, would be disrupted. We analyzed the expression of Megf10, a CAM involved in ChAT^+^ amacrine cell spacing (Kay et al., 2012) and Fat3, a cadherin involved in dendrite formation and migration of ChAT^+^ amacrine cells (Deans et al., 2011). In P14 wild-type retinas, Megf10 expression overlapped with ChAT in the cell soma and in the IPL (Figure 4A). In P14 *Pten*^cKO^ retinas, expression of Megf10 in ChAT^+^ soma was comparable to wild-type retinas, but in the IPL, Megf10 expression was severely disturbed, with large patches of Megf10 staining that did not co-localize with ChAT (Figure 4B). Similarly, Fat3 is normally co-localized with ChAT in IPL processes and not in the soma in P14 wild-type retinas (Figure 4C). In contrast, in P14 *Pten*^cKO^ retinas, Fat3 localization was strikingly disrupted, and similar to Megf10, large Fat3 expressing patches were observed in the IPL and in the INL that did not co-localize with ChAT staining (Figure 4D).

**Figure 4.**
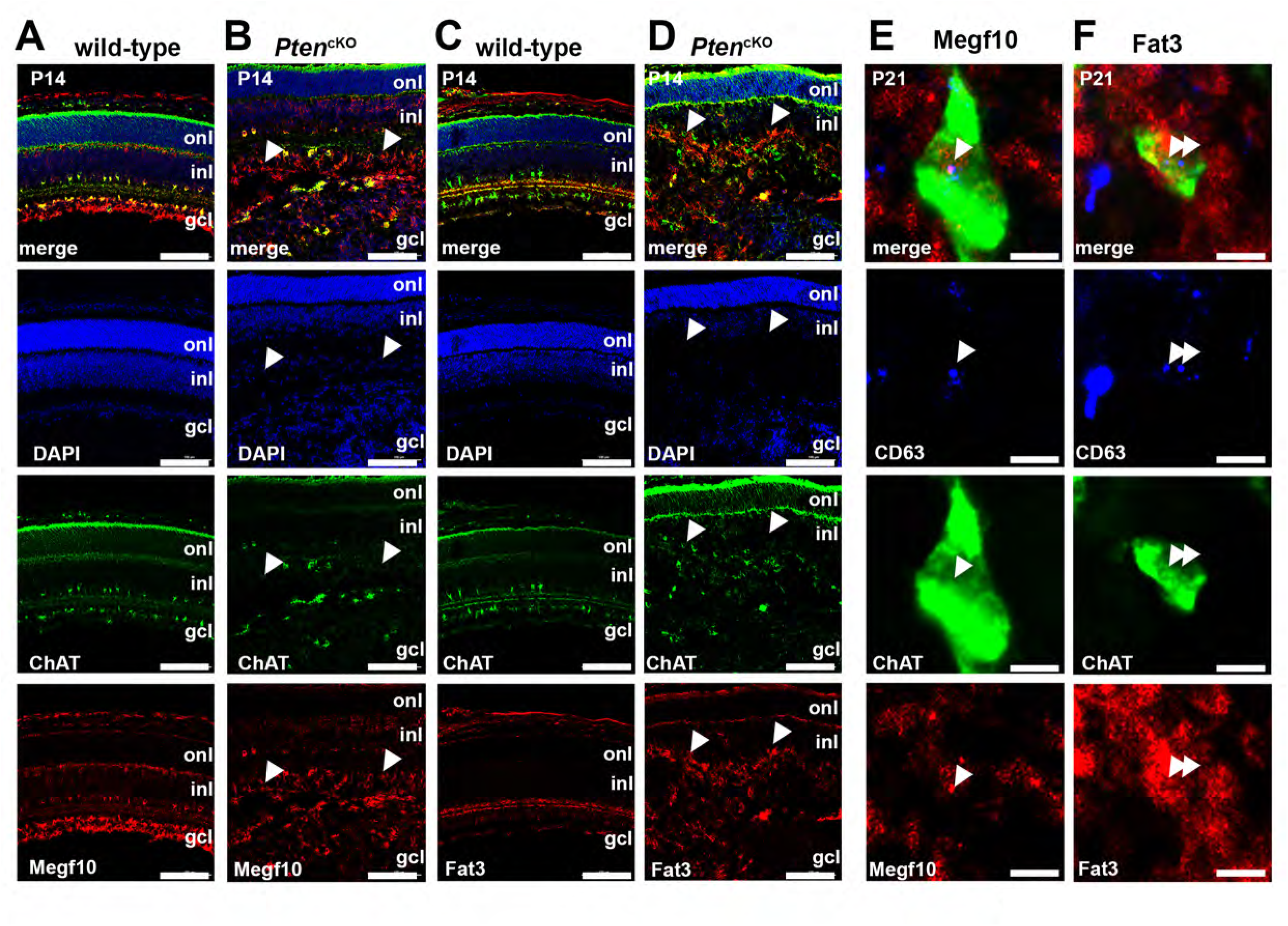
Co-localization of Megf10 and Fat3 with ChAT is disrupted in *Pten*^cKO^ retinas. (A-B) Colabeling of P21 wild-type (A) and *Pten*^cKO^ (B) retinas with ChAT (green) and Megf10 (red). Blue is DAPI counterstain of nuclei. Arrows in B show abnormal expression of Megf10 in the absence of ChAT in the *Pten*^cKO^ IPL. (C-D) Colabeling of P21 wild-type (C) and *Pten*^cKO^ (D) retinas with ChAT (green) and Fat3 (red). Blue is DAPI counterstain of nuclei. Arrows in D show abnormal expression of Fat3 in the absence of ChAT in the *Pten*^cKO^ IPL. (E,F) Colabeling of P21 wild-type retinas with CD63 (blue), ChAT (green) and Megf10 (red) (E). Arrows point to endosomes. Colabeling of P21 wild-type retinas with CD63 (blue), ChAT (green) and Fat3 (red) (F). Arrows point to endosomes. gcl, ganglion cell layer; inl, inner nuclear layer; onl, outer nuclear layer. Scale Bar: A-D: 100μm, E-F: 10μm.

Next, we asked whether Megf10 and Fat3 were present in endosomes, as expected for CAMs, by examining co-expression with CD63, a marker of multivesicular bodies (MVBs)/late endosomes/lysosomes (Kobayashi et al., 2000). As expected, both Megf10 and Fat3 co-localized with CD63 in ChAT^+^ amacrine cells (Figure 4E,F), consistent with the general idea that CAMs are normally remodeled through recycling endosomes (Kawauchi, 2012). As a corollary, the disruption of Megf10 and Fat3 expression in *Pten* cKO retinas (Figure 4A-D) indicates that CAM remodeling is disrupted in the absence of this intracellular phosphatase.

### Pten regulates Dscam localization in dopaminergic amacrine cells

As previously shown (Cantrup et al., 2012), TH^+^ amacrine cell somal mosaics and their associated dendritic nets are also disrupted in *Pten*^cKO^ retinas (Figure 5A). A similar phenotype is observed in *Dscam*^del17^ mutants, although the fasciculation of TH^+^ dendrites was more severe (Figure 5A), consistent with previous reports (Fuerst et al., 2009; Fuerst and Burgess, 2009; Fuerst et al., 2010; Fuerst et al., 2008; Keeley et al., 2012). We therefore questioned whether the endocytic localization of Dscam was disrupted in *Pten*^cKO^ TH^+^ cells. For this purpose, we assessed the distribution of Dscam in TH^+^ amacrine cells in *Pten*^cKO^ retinas, which has been documented to increasingly acquire a more punctate pattern beginning at P2 (de Andrade et al., 2014). Notably, Dscam is also expressed on the cell membrane, which can be detected with other antibodies (Li et al., 2015), but our interest was specifically in the vesicular compartment carrying this protein. In P21 wild-type retinas, over 98% of TH^+^ amacrine cells had 1-3 Dscam^+^ puncta, with only a few cells (1 .44 %) having more than 3 puncta (Figure 5B,C). In striking contrast, in P21 *Pten*^cKO^ retinas, over 40% of TH^+^ amacrine cells contained more than 3 Dscam^+^ puncta, with up to 7-9 puncta per cell, which was never observed in wild-type retinas (Figure 5B,C). The P21 *Pten*^cKO^ distribution thus clearly shifted to higher numbers of Dscam^+^ puncta per cell, and was confirmed to be significantly different (G test: p=0.013) (Figure 5C).

**Figure 5.**
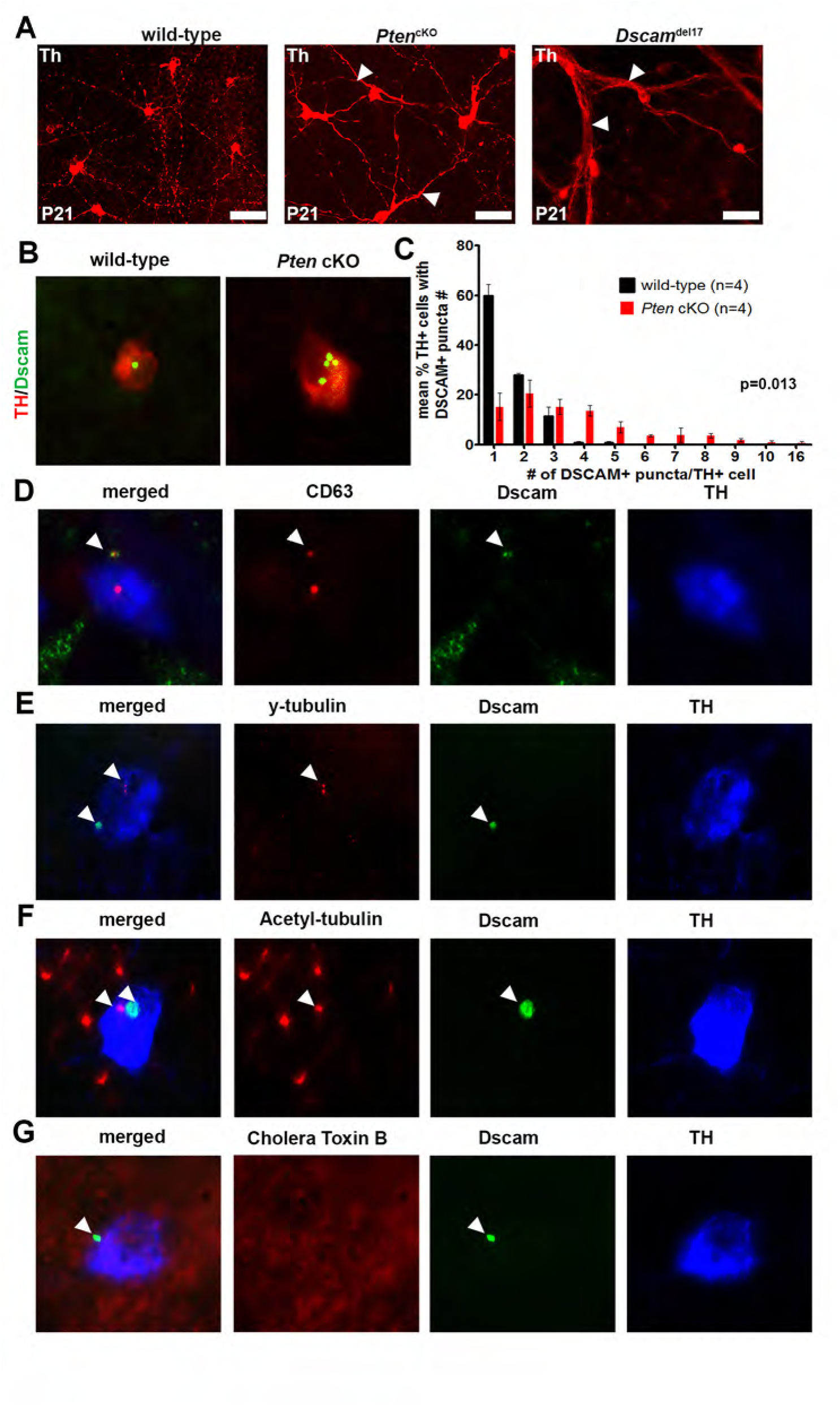
Dscam accumulates aberrantly in TH+ dopaminergic amacrine cells in *Pten*^cKO^. (A) Immunolabeling of P21 wild-type, *Pten*^cKO^ and *Dscam*^del17^ retinal flatmounts with TH. Arrowheads point to TH^+^ amacrine cell process fasciculation. (B) Colabeling of P21 wild-type and *Pten*^cKO^ retinas with TH (red) and Dscam (green). (C) Quantification of the number of Dscam^+^ puncta localizing within individual TH^+^ cells in P21 wild-type and *Pten*^cKO^. (D-G) Colabeling of P21 wild-type retinas with DSCAM (red) and TH (green) along with the following cell type markers: CD63 (D), γ-tubulin (E), acetyl-tubulin (F) and cholera toxin B (G). G test = p<0.05, indicating the distributions are significantly different.

Next, we asked whether Dscam^+^ puncta localized to specific subcellular organelle(s) by performing triple immunolabelling with TH, Dscam, and organelle markers, including CD63, to mark MVBs/late endosomes/lysosomes (Figure 5D) γ-tubulin, to mark centrosomes (Figure 5E), acetylated tubulin, to mark the primary cilium (Figure 5F), and cholera toxin β, to mark lipid rafts (Figure 5G). Of these markers, Dscam puncta co-localized with CD63 in TH^+^ amacrine cells, indicating that Dscam protein largely forms puncta, localizing to MVBs, late endosomes and/or lysosomes (Figure 5D).

Taken together, these data suggest that *Pten* controls the amount of Dscam protein sequestered into recycling endosomes in TH^+^ amacrine cells, which may prevent its normal recycling to the cell membrane.

### Retinal cells secrete extracellular vesicles

Late endosomes, or MVBs, are comprised of multiple intraluminal vesicles that are packed with proteins, lipids, and nucleic acids, which can be trafficked to lysosomes for degradation, or to the plasma membrane, where they undergo exocytosis and are released as EVs (Figure 6A). In the eye, EVs are secreted from the retinal pigmented epithelium (RPE) and corneal fibroblasts, and are found in the aqueous humor and in tears, but little is known about EV secretion from the retina itself (Klingeborn et al., 2017). To determine whether retinal cells secrete EVs, the RPE was removed from P21 retinas, which were then dissociated and plated for 24 hr, followed by conditioned media (CM) collection and EV isolation using differential centrifugation (Colombo et al., 2014). Using transmission electron microscopy (TEM), a large number of membrane-enclosed vesicles in the predicted size range of 50-150 nm for small EVs (or exosomes) were detected in P21 retinal cell CM (Figure 6B). To confirm that these vesicular-like structures were indeed EVs, they were immunolabeled with CD9, an EV-specific tetraspanin, and their size distribution was monitored via nanoscale-flow cytometry, confirming the presence of CD9^+^ particles (Figure 6C).

**Figure 6.**
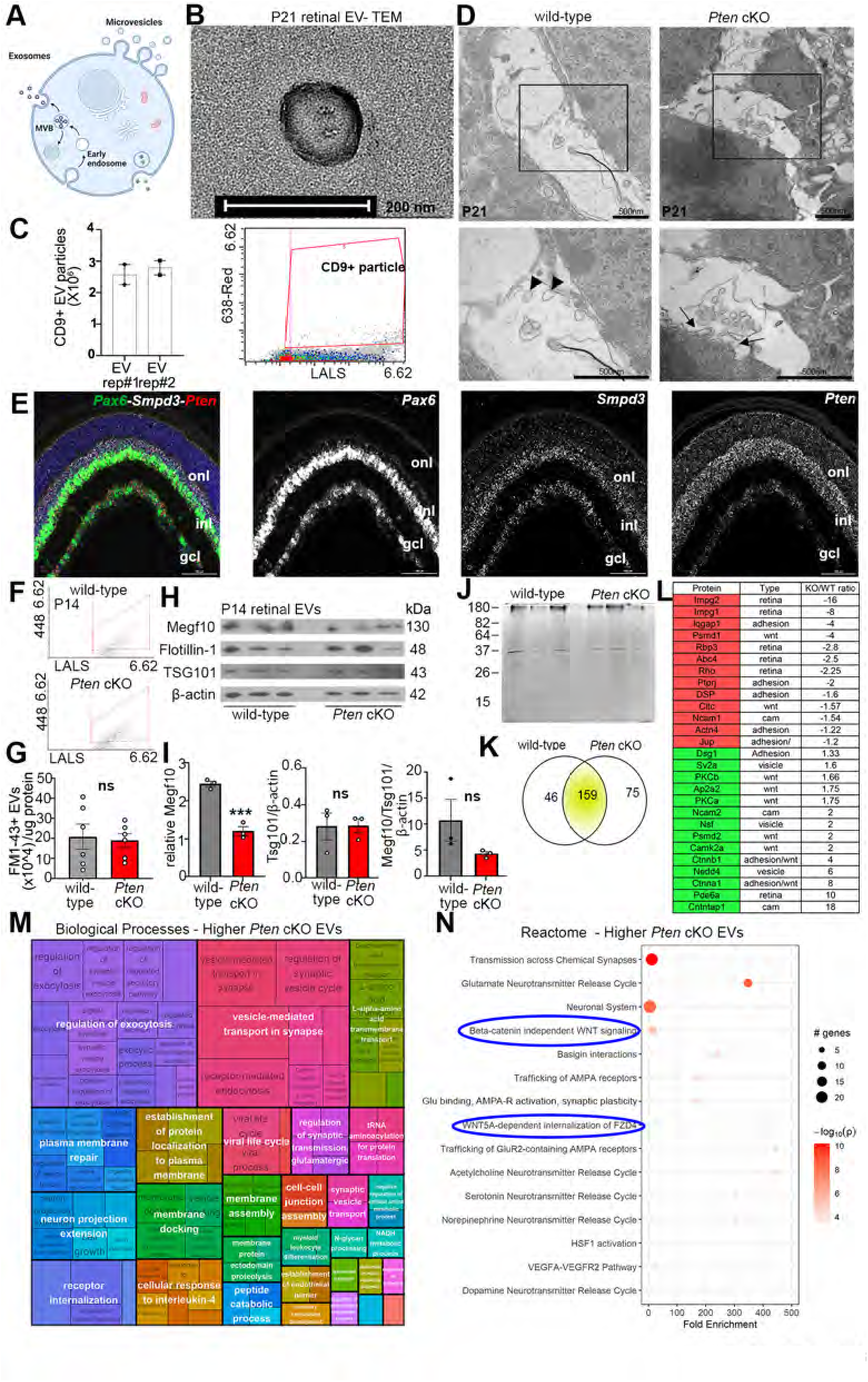
Retinal cells secrete extracellular vesicles containing adhesion molecules known to regulate amacrine cell spacing. (A) Schematic of retinal cell showing endosomes, multi-vesicular bodies (MVB) and secreted exosomes. (B) Transmission Electron Microscopy (TEM) of an exosome purified from P12 wild-type retina. (C) Nanoflow analysis of CD9-labeled EVs from P12 retina. (D) TEM of P21 wild-type and *Pten*^cKO^ retinas focusing on the lower INL, where amacrine cells reside. Boxed areas are magnified in panels below and show membrane enclosed vesicles (arrowheads), including a MVB in the process of being pinched off and extruded from the cell (arrows). (E) RNAscope analysis of *Pax6* (green)*, Smpd3* (white) and *Pten* (red) expression in P21 retinas. Blue is DAPI counterstain. (F,G) Nanoflow quantification of EVs extracted form P14 wild-type and *Pten*^cKO^ retinas. (H) Western blot analyses and densitometry for Megf10 and EV markers Flotillin-1 and TSG101 and β-actin on EVs isolated from P14 wild-type and *Pten*^cKO^ retinal EVs. (I) Silver stained SDS-PAGE gel with EV lysates purified from P14 wild-type and *Pten*^cKO^ retinas. (J) LC-MS/MS analyses of EV proteins isolated from P14 wild-type and *Pten*^cKO^ retinas, showing the number of proteins enriched in wild-type and *Pten*^cKO^ samples. (K) A list of differentially enriched proteins in P14 wild-type and *Pten*^cKO^ retinal EVs. Red shaded proteins in H indicates higher level in *Pten*^cKO^, whereas green shaded proteins are lower in *Pten*^cKO^ retinal EVs. (L) GO biological processes and KEGG pathways analysis of enriched proteins in *Pten*^cKO^ retinal EVs. (M) Reactome pathway analysis of enriched proteins in *Pten*^cKO^ retinal EVs. The size of the circle represents the number of genes enriched in the pathway; the color of the circle represents the log10 P-value. p values are denoted as follow: <0.05 *. Scale Bars: B: 200nm; D: 500nm; E: 100μm; See also Figure S4.

To further characterise these vesicles *in situ*, we used TEM and observed membrane-enclosed, vesicular-like structures secreted in the extracellular space in the inner INL of P21 retinas, where amacrine cells reside (Figure 6D). Secreted vesicles were observed in P21 wild-type and *Pten*^cKO^ retinas, including individual vesicles that appeared to be attaching to the cell membrane (Figure 6D), and larger clusters of vesicles enclosed in a membrane that appeared to be a multivesicular cargo just emerging from the plasma membrane, as shown in a P21 *Pten*^cKO^ retina (Figure 6D). Strikingly, the sizes of the observed vesicles were all around 50-90 nm in diameter, which corresponds to the size range of exosomes (50-150 nm; reviewed in (Thery et al., 2002). Of note, these membrane-bound structures are not synaptic vesicles, as synaptic vesicles are much smaller, ranging from 20-40 nm with a median size of 30 nm for bipolar cells (Matthews and Sterling, 2008). We thus have evidence that EVs, and more specifically, exosomes, are secreted by retinal cells.

### Smpd3-mediated EV production is not required for amacrine cell spacing

As our observations suggested that the EVs secreted from retinal cells were in the exosome size range, we next assessed the expression of the EVs pathway machinery in the retina. We focused on neutral sphingomyelinase 2 (nSmase2), encoded by *Smpd3,* which hydrolyses sphingomyelin to ceramide, a necessary first step in exosome production (Kosaka et al., 2010). Strikingly, using RNAscope in situ hybridization, *Smpd3* transcripts were largely restricted to the inner INL and the GCL, closely matching the pattern of *Pax6* transcript distribution, an amacrine cell marker, and overlapping with *Pten*, which was expressed throughout the INL (Figure 6E). To test the functional significance upon amacrine cell spacing in the absence of nSmase2, we acquired *fro/fro* mice, which harbor a spontaneous mutation in *Smpd3*, the gene encoding for this enzyme (Alebrahim et al., 2014). Interestingly, both ChAT^+^ and TH^+^ amacrine cell mosaics were normal in adult *Smpd3* null mutant mice (Figure S4A,B). While this data might suggest that *Smpd3* does not control amacrine cell spacing, we reasoned that other ESCRT pathway components may compensate for the loss of *Smpd3*. Thus, we focused instead on investigating whether Pten regulated retinal EV number or content.

### *Pten* loss alters vesicle mediated transport and the EV proteome

We first asked whether *Pten* controls retinal EV secretion by quantifying the number of EVs secreted in *Pten*^cKO^ retinas using nanoscale-flow cytometry, labeling EVs with FM1-43, a membrane dye. Compared to P14 wild-type retinas, the relative number of EVs secreted in *Pten*^cKO^ retinas, normalized to the protein content of input cells, was not significantly different (Figure 6F,G). However, even though the number of EVs was not affected in *Pten*^cKO^ retinas, we reasoned that *Pten* could still be involved in allocating the cargo packaged into the EVs. We first tested this idea by western blotting, focusing on Megf10, a CAM that regulates amacrine cell spacing (Kay et al., 2012). We focused on Megf10 as it has an altered distribution in P14 *Pten*^cKO^ ChAT^+^ amacrine cells (Figure 4), suggesting that it may be secreted into the extracellular space via EVs. Retinal EVs were isolated via differential centrifugation from P14 wild-type and *Pten*^cKO^ retinas, the presence of which we validated by examining the expression of EV-specific markers, including flotillin and TSG101 (Colombo et al., 2014) (Figure 6H). The absolute amount of Megf10 was significantly reduced in P14 *Pten*^cKO^ retinal EVs (n=3), and there was a trend towards reduced levels normalized to EV quantity, even though vesicular content, as monitored by TSG101 levels, did not differ compared to wild-type retinas (Figure 6I). These data indicate that in the absence of *Pten*, EV cargo changes, prompting us to ask which other proteins might be secreted in retinal EVs.

We next performed an unbiased assessment of retinal EV proteins, comparing the content of EVs isolated from P14 wild-type and *Pten*^cKO^ retinas by liquid chromatography-tandem MS (LC-MS/MS) (n=3 each; Figure 6J). A total of 280 proteins were detected in P14 EVs. 159 were present in both wild-type and *Pten*^cKO^ mutant retinas, with 46 unique wild-type and 75 unique *Pten*^cKO^ EV proteins (Figure 6K). Proteins that were expressed at equivalent levels in both samples included several ER/Golgi proteins that are also secreted in EVs (Emc1, Glg1, Atp2a2) (Figure 6L). Higher abundance proteins in the wild-type retina included several proteins expressed in photoreceptors, including Abca4, Rho, Rbp3, Impg1 and Impg2 (Figure 6L). The reduction in photoreceptor protein expression in *Pten*^cKO^ retinas was consistent with our previous report of fewer rod photoreceptors in these mutant mice (Tachibana et al., 2016). Conversely, several vesicle-related proteins were exclusively found in *Pten*^cKO^ retinas, including Nedd4, which promotes the export of PTEN into EVs (Putz et al., 2012), Contactin Associated Protein 1 (CNTP1), which is a transmembrane protein expressed in amacrine cells of unknown function (O’Brien et al., 2010) and Pde6a, the mutation of which is associated with Retinitis Pigmentosa and photoreceptor degeneration (Mowat et al., 2017) (Figure 6L). In addition, other vesicle and ER proteins (Nsf, Sv2a, Hsp90b1, Erap1) were enriched in *Pten*^cKO^ EVs. Finally, two cell adhesion molecules that were differentially expressed – Ncam2, which was enriched in *Pten*^cKO^ EVs, is associated with neurodevelopmental disorders (Petit et al., 2015) and neurite branching (Sheng et al., 2015), and Ncam1, which was downregulated in *Pten*^cKO^ EVs, regulates amacrine cell neurite growth (Kljavin et al., 1994), the disruption of which influences amacrine cell mosaic order (Figure 6L).

To better understand how *Pten* affects EV content, we performed a global gene ontology (GO) analysis of the proteomic data collected from the EV cargo (Figure 6M; Table S1). The biological processes enriched in P14 *Pten*^cKO^ retinal EVs included “*regulation of exocytosis*”, “*vesicle-mediated transport in synapse*”, “*plasma membrane repair*”, “*establishment of protein localization to plasma membrane*”, and “*membrane assembly*” (Figure 6M; Table S3). These categories differed from those enriched in P14 wild-type retinal EVs, which were mainly associated with ion transport (Table S2). In addition, Reactome analysis of signaling pathways identified “*beta-catenin independent Wnt signaling”, Wnt5a-dependent internalization of Fzd4*”) as enriched in P14 *Pten*^cKO^ retinal EVs (Figure 6N; Table S5), whereas ion transport and ion homestasis predominated in P14 wild-type retinal EVs (Table S4). Enriched proteins in P14 *Pten*^cKO^ retinal EVs included α-Catenin, β-Catenin, PKCα, CAMK2a and Psmd2, all involved in canonical and non canonical Wnt signalling, prompting us to further assess the role of Wnt signaling in amacrine cell distribution.

### *Pten*-dependent canonical Wnt signalling controls the spatial distribution of ChAT^+^ amacrine cells

Laminar attractants are thought to guide migrating amacrine cells from the apical surface of the developing retina to the future INL/GCL, but their identity is unknown (Galli-Resta, 2002). From our Reactome analyses of EVs, Wnt signaling appeared to be altered in *Pten*^cKO^ retinas. This result was not surprising given our finding that *Pten* regulates endosomal trafficking, and Wnts are large hydrophobic molecules that are secreted in EVs in order to mediate long-range signaling (Gross et al., 2012). Noncanonical Wnt signaling has been assessed in the retina and shown to be important for the development of the outer plexiform layer (Sarin et al., 2018) whereas canonical Wnt signaling in the developing retina is understudied (Figure 7A). Canonical Wnts, such as Wnt3a and Wnt2b, bind to frizzled ligands to ultimately stabilize β-catenin and allow its translocation into the nucleus, where β-catenin binds LEF/TCF transcription factors to activate downstream target genes (Figure 7A). *Wnt2b* and *Wnt3a* transcripts were detected at the highest levels in the developing INL and GCL in P0 wild-type and *Pten*^cKO^ retinas (Figure 7B). To determine which cell types display active Wnt signaling in the retina, we used a *lef/tcf-lacZ* reporter (Liu et al., 2003). β-galactosidase (β-gal) immunostaining was detected in the developing inner INL and GCL of P4 wild-type retinas, where amacrine cells are located, co-localizing with Pax6, an amacrine cell marker (Figure 7C). To measure levels of Wnt signaling, we monitored non-phosphorylated (i.e., active) β-catenin expression, which was detected at low levels in the IPL of P14 wild-type retinas, and sharply upregulated in *Pten*^cKO^ retinas, both by immunostaining (Figure 7D) and by western blotting (Figure 7E,F). Canonical Wnt signalling is thus active in migrating amacrine cells, and disturbed in *Pten*^cKO^ retinas.

**Figure 7.**
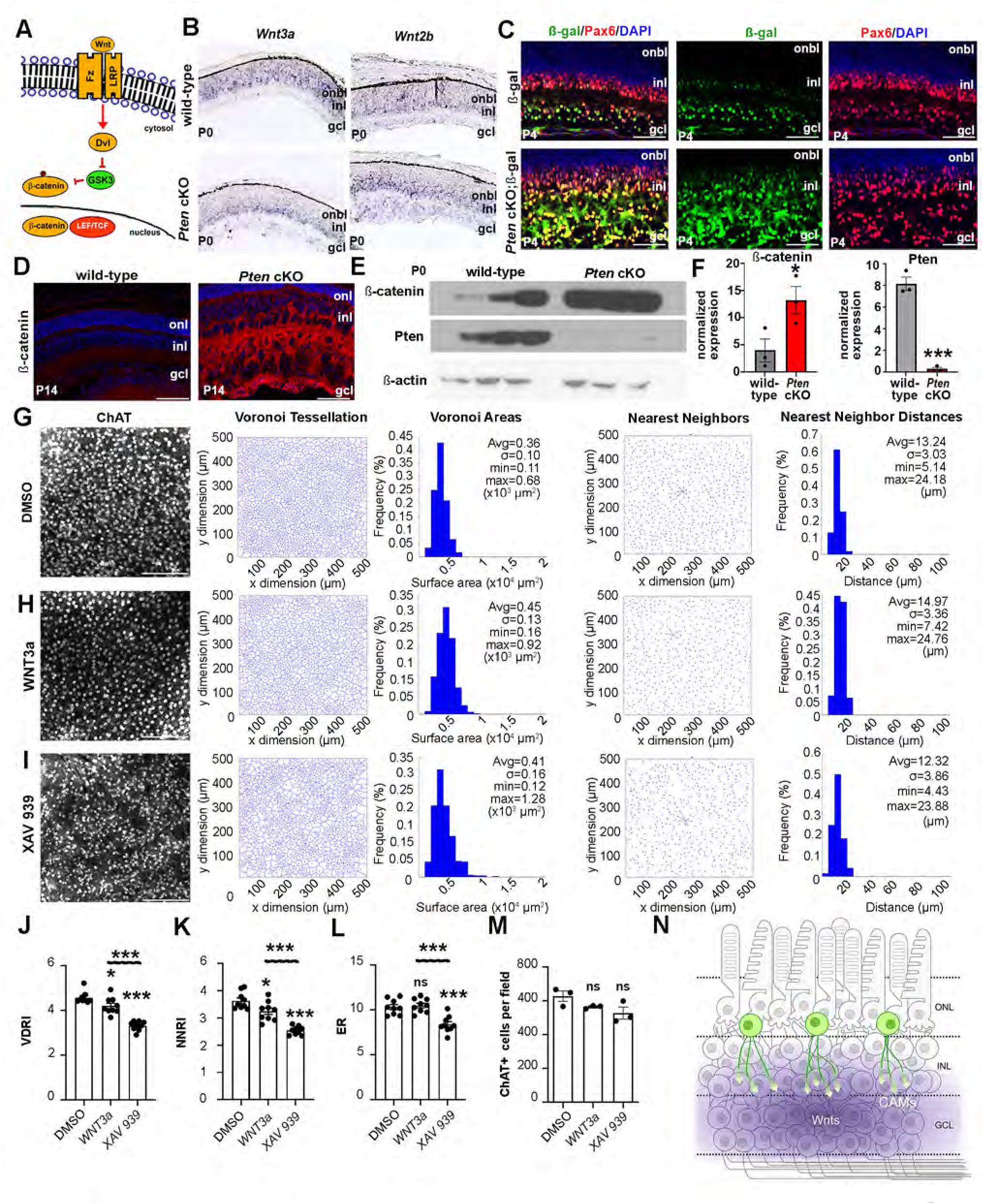
Wnt signalling pathway alteration perturbs cholinergic amacrine cell tangential spatial distribution. (A) Schematic illustration of canonical Wnt signalling pathway. (B) Expression of *Wnt3a* and *Wnt2b* in P0 wild-type and *Pten*^cKO^ retinas using RNA *in situ* hybridization. (C) Expression of β-galactosidase (β-gal) and Pax6 in P4 *lef/tcf::lacz* and *Pten*^cKO^; *lef/tcf::lacz* retinas. (D) Expression of β-catenin in P14 wild-type and *Pten*^cKO^ retinas. (E,F) Western blot analysis of β-catenin and Pten expression levels in P0 wild-type and *Pten*^cKO^ retinas, normalized to β-actin using densitometry. (G-I) ChAT^+^ immunolabeling and mosaic analysis of wild-type retinal explants cultured with DMSO (G), Wnt3a (H) and XAV939 (I). Voronoi Tessellation of ChAT^+^ amacrine cells are shown, as well as nearest neighbor plots and frequency distributions of nearest neighbor distances between ChAT^+^ amacrine cells. (J-M) Calculation of Voronoi domain (J) and nearest neighbor (K) regularity indices, effective radius (L) and ChAT^+^ cell number per field (M) in DMSO, Wnt3a and XAV939 treated retinal explants. (N) Schematic illustration of model of amacrine cell migration, with Wnt activation and endocytic remodeling of CAMs playing a critical role in this process. p values are denoted as follow: <0.05 *, <0.01 **, <0.001 ***. onbl, outer neuroblast layer. Scale Bars: C: 50μm; D: 100μm; G-I: 200μm.

To address whether canonical Wnt mediates the spatial distribution of amacrine cells downstream of *Pten*, we used small molecules to activate or inactivate Wnt signalling and analyzed effects on the tangential distribution of ChAT^+^ amacrine cells. P0 retinal explants were cultured for 7 days *in vitro* (DIV), during the period of active amacrine cell migration, with either DMSO (control; Figure 7G), recombinant Wnt3a ligand (Figure 7H) or XAV939 (Figure 7I), a tankyrase inhibitor that increases the axin-GSK3β destruction complex that degrades β-catenin (i.e., Wnt inhibitor) (Huang et al., 2009). After 7DIV, explants were immuno-stained with ChAT and the spatial distribution of ChAT^+^ amacrine cells was assessed using Voronoi domain and nearest neighbor analyses. In all conditions, ChAT^+^ amacrine cells were very densely packed (Figure 7G-I), not respecting the ∼ 3 diameter exclusion zones seen in normal wild-type retinas, presumably a consequence of reduced retinal expansion *in vitro*. In explants cultured with Wnt3a (Figure 7H) and especially for XAV939 (Figure 7I), ChAT^+^ cells appeared less regularly distributed relative to the DMSO (control) treated explants, as revealed by calculations of VDRI (Figure 7J) and NNRI (Figure 7K). We also calculated the effective radius (ER) which indicates the territory surrounding each cell where no other like-type cells can be found, averaged across the entire population of cells (Figure 7K). The ER was significantly reduced in the XAV939-treated explants compared to both DMSO control and Wnt3a treated explants (n=3; p=0.0079; Figure 7L), indicating that many cells in this condition are positioned a little closer to one another than in the other two conditions. Notably, occasional gaps in the mosaic were seen in XAV939-treated explants, a further indication of patterning defects (Figure 7I). Notably, these spacing alterations arose without impacting the overall number of ChAT^+^ cells between the groups (Figure 7M).

Taken together, these data indicate that blocking Wnt signalling with XAV939 partially phenocopies the perturbation of ChAT^+^ amacrine cells observed in *Pten*^cKO^ retinas. Our model is that Wnt ligand expression in the lower retina attracts and activates canonical signaling in migrating amacrine cells (Figure 7N). The full extent of the aberrant amacrine cell distribution in *Pten*^cKO^ retinas is thus likely arising both from altered signaling through the Wnt pathway, and the aberrant endocytic recycling of CAMs on the cell surface.

## DISCUSSION

In 1942, Waddington proposed the concept of developmental canalization, which suggests that built-in constraints ensure that optimal developmental programs are followed to ensure constancy of form. However, the molecular nature of the built-in constraints that pattern most tissues remains incompletely understood. In this article, we asked how amacrine cells, which form regular mosaics in the retina, are patterned at the molecular level. Previously, *Pten* (Cantrup et al., 2012) and *Dscam* (Fuerst et al., 2009; Fuerst et al., 2008) were shown to regulate the radial and tangential positioning of TH^+^ amacrine cells as well as their dendritic fasciculation. However, from these early studies, it was suggested that Dscam was not expressed in ChAT^+^ amacrine cells. By applying new single-cell transcriptomic and RNAscope technologies, we were able to show that *Pten* and *Dscam* transcripts are indeed co-localized in ChAT^+^ amacrine cells. Furthermore, by performing single and double mutant analyses, we revealed that *Pten* and *Dscam* act in parallel, partially redundant pathways to regulate tissue patterning, such that the most severe patterning disruptions are seen in *Pten;Dscam* double mutants. At the genetic level, these studies suggest that *Pten* and *Dscam* act in parallel to regulate amacrine cell mosaic spacing, but whereas *Pten* and *Dscam* are both essential contributors to the formation of TH^+^ mosaics, ChAT^+^ mosaics primarily depends on *Pten,* with a minor contribution from *Dscam*, as *Dscam* loss-of-function affected ChAT^+^ spacing only in the context of a *Pten*^cKO^.

At the molecular level, we found that *Pten* regulates endosomal recycling of Dscam, Megf10, and Fat3, three CAMs involved in amacrine cell spacing, indicating that the Pten and Dscam pathways are not entirely parallel. Using a proteomic screen, we identified a general role for *Pten* in controlling vesicle-mediated trafficking, including of Wnt signaling molecules, which are large hydrophobic molecules that are packaged in extracellular vesicles for long-range signaling (Gross et al., 2012). Wnt signaling had not previously been implicated in amacrine cell spacing, but we show that Wnt signaling is activated in migrating amacrine cells, and its pharmacological perturbation phenocopies *Pten*^cKO^ amacrine cell patterning defects. We further found that *Pten* normally inhibits Wnt signaling, as expected, because in the absence of *Pten*, PI3K/Akt signaling is elevated in the retina (Tachibana et al., 2016), and Akt inhibits GSK3, a central component of the β-catenin destruction complex (Tanaka et al., 2011). Consequently, β-catenin is stabilized and Wnt signaling is elevated in *Pten*^cKO^ retinas. Taken together, these data reveal new insights into the mystery of how an intracellular phosphatase regulates the patterning of cells that rely on cell surface interactions mediated by cell adhesion molecules, as well as a Wnt gradient. They further position *Pten*-regulated endocytic trafficking as a novel constraint mechanism to control retinal cell patterning.

Previously, we revealed that amacrine cells are mispositioned in the radial plane in *Pten*^cKO^ retinas (Cantrup et al., 2012). These defects in radial migration are not a secondary consequence of the reduction in amacrine cell number previously documented in *Pten*^cKO^ retinas (Tachibana et al., 2016). Indeed, amacrine cell differentiation and migration are separable events. For example, increasing amacrine cell number with a Tgfβ2 antagonist does not influence radial patterning (Ma et al., 2007). Moreover, retinal expansion is also observed in *p27*/*Cdkn1b* KOs, one of the downstream targets of PI3K/PTEN signaling (Mamillapalli et al., 2001; Tokita-Ishikawa et al., 2010). However, unlike in *Pten* or *Dscam* mutants, in *Cdkn1b* KOs each layer is expanded, and there is no mispositioning of retinal cells in the IPL. Moreover, *Pten* mutants have fewer cells overall, so a hyperproliferative phenotype does not underlie the expansion of the IPL (Tachibana et al., 2016).

*Dscam* KOs (Fuerst et al., 2009; Fuerst et al., 2008) have very similar defects in the radial migration of amacrine cells as *Pten*^cKO^ retinas. Through the analysis of double *Pten*;*Dscam* mutants we found that *Dscam* and *Pten* function in parallel pathways to regulate INL and GCL lamination. How Dscam regulates the radial migration of amacrine cells is unknown, but it is likely related to its ability to act as a ‘non-stick coating’, which prevents cells from clumping together (Fuerst et al., 2012; Fuerst et al., 2009; Fuerst and Burgess, 2009). With respect to *Pten*, the most likely explanation for the defects in amacrine cell migration are that mutant amacrine cells are not properly coated with adhesion molecules. Indeed, defects in CAM endocytic recycling were evident when examining the expression of amacrine cell CAMs, such as Fat3, which ectopically localized to amacrine cell bodies in *Pten*^cKO^ (this study), suggesting that the multipolar→ bipolar→ unipolar transitions may be perturbed (Ellenbroek et al., 2012). In addition, other CAMs that act as homophilic repellants, such as *Megf10* (Kay et al., 2012), were also aberrantly expressed in Pten cKO retinas. Intriguingly, in gain-of-function assays, *Megf10* non-cell autonomously influences amacrine cell mosaics (Kay et al., 2012). These data are consistent with the idea that secreted repellants may control amacrine cell mosaics, which has been proposed, but not yet proven (Galli-Resta et al., 2008).

EVs were initially implicated as a waste management system to get rid of unwanted cellular components, but they are now viewed as an essential mode of intercellular signaling. We found here that Dscam and Megf10 co-localize with CD63 in retinal amacrine cells, a tetraspanin that is highly expressed in late endosome, lysosome, and MVBs (Kanuma et al., 2017), but are also part of EVs. Interestingly, a previous study revealed that the N-terminal region of Dscam can be secreted into the media *in vitro*, but potential functions in neural development had not been identified, and it was not shown whether Dscam was secreted in EVs (Schramm et al., 2012). While we did not directly show that the retinal EVs we isolated were produced by amacrine cells, based on EM, there appear to be MVBs secreted into the extracellular space in the inner INL, where amacrine cells reside. We have characterized the contents of these EVs, and identified Ncam1 and Ncam2 as additional cell adhesion molecules that were differentially expressed in wild-type and *Pten*^cKO^ retinal EVs. Future studies will test the significance of these CAMs with respect to amacrine cell spacing. Finally, Wnt signalling pathway proteins were differentially enriched in *Pten*^cKO^ retinal EVs, and when we perturbed Wnt signalling, the spatial organization of cholinergic amacrine cells was disrupted, indicating an important role of Wnt in this process.

In summary, through our analyses of *Pten* and *Dscam* interactions, we have revealed that two independent pathways regulate amacrine cell spacing in both the radial and tangential axis, and furthermore, we have found that endocytic trafficking of CAM and Wnt proteins, and their secretion into EVs, contributes to the process by which normal amacrine cell spacing is achieved.

## ACKNOWLEDGEMENTS

We thank Natasha Klenin (University of Calgary) and Dawn Zinyk and Lakshmy Vasan (Sunnybrook Research Institute) in the Schuurmans’ lab for technical assistance, Dennis Lee in Dr. Hon Leong’s lab (Sunnybrook Research Institute) for reagents and technical support, and Sarah Dalesman (University of Calgary) for assistance with statistical analyses of Dscam^+^ puncta. We also thank Jung-Lynn (Jonathan) Yang for assistance with artwork (University of Calgary). This work was supported by the Canadian Institutes of Health Research (CIHR) MOP-142338 and CIHR PJT 180243 to CS, YT and IK, and by the National Institutes of Health (NIH) to BER (EY-019968). JH is supported by a Canada Graduate Scholarship – Doctoral (CGS-D)/CIHR award, Vision Science Research Program Scholarship, Ontario Graduate Scholarship and R.O. Torrance Bursary. NT and RC were supported by studentships from the ACHRI/CIHR Training Grant at the University of Calgary, and by a Lion Sight Center Award. CS holds the Dixon Family Chair in Ophthalmology Research.

## AUTHOR CONTRIBUTION

YT,JH,NT: conceptualization, data curation, formal analysis, investigation, methodology, visualization, validation, writing – original draft, writing – review and editing

LAD,SO,TO,YI,VC,FS,AB,RD,RC: formal analysis, investigation, methodology, validation, writing – review and editing

PM,PEM,JLL: investigation, methodology, validation, writing – review and editing

MM,MC,IK,MB,AdS: resources, supervision, writing – review and editing

BER: funding acquisition, conceptualization, formal analysis, methodology, writing – review and editing

CS:project administration, resources, supervision, validation, writing – original draft; writing – review and editing

## STAR METHODS

### Resource Availability

#### Lead Contact

Further information and request for resources and reagents should be directed to and will be fulfilled by the Lead Contact, Dr. Carol Schuurmans.

#### Materials Availability

Transgenic mouse lines generated in this study are deposited in a central repository (Jackson Laboratory). All antibodies and reagents are commercially available.

#### Data and Code Availability

The mass spectrometry proteomics data have been deposited to the ProteomeXchange Consortium via the PRIDE [1] partner repository with the dataset identifier PXD036221 and 10.6019/PXD036221. Uncropped western blot images were deposited in Mendeley data (Mendeley Data, V1, doi: 10.17632/brd87jgd82.1). Accession numbers are listed in the Key Resources Table. These data are publicly available as of the date of publication. All other data reported in this study will be shared by the lead contact upon request.

### Experimental Models and Subject Details

None of our experimental animals had been previously used for other procedures. All animals were healthy.

#### Mice

All animal procedures were approved by the University of Calgary Animal Care Committee (AC11-0053) and later by the Sunnybrook Research Institute Animal Care Committee (16-606) in agreement with the Guidelines of the Canadian Council of Animal Care (CCAC). The following animal lines were used from Jackson Laboratory: *Pten^fl^* allele (Backman et al., 2001) (B6.129S4-*Pten^tm1Hwu^*/J. Strain #: 006440, RRID:IMSR_JAX:006440), *Pax6::Cre* driver (Marquardt et al., 2001) (STOCK Tg(Pax6-GFP/cre)1Rilm/J. Strain #:024578. RRID:IMSR_JAX:024578. Common Name: P0-3.9GFPCre), *fro/fro* (Alebrahim et al., 2014), *Dscam^del17^* (Fuerst et al., 2008) B6.CBy-*Dscam^del17^*/RwbJ. Strain #:008000. RRID:IMSR_JAX:008000) and lef/tcf-lacZ reporter mice (STOCK Tg(TCF/Lef1-lacZ)34Efu/JStrain #:004623. RRID:IMSR_JAX:004623. Common Name: TOPGAL). PCR genotyping was performed as described by Jax. Lines were maintained by crossing with C57BL/6J wild-type mice (Strain #:000664. RRID:IMSR_JAX:000664. Common Name: B6). Animal explants were performed with CD1 mice (Charles River, IMSR_CRL:022). The day of the vaginal plug was considered to be embryonic day (E) 0.5 for timed pregnancies.

### Method Details

#### Tissue processing

Eyes were dissected out from embryos or postnatal pups at the designated stages and either fixed directly or the retinas were first dissected out. Fixation of eyes or retinas was achieved by immersing tissues in 4% paraformaldehyde (PFA)/1X phosphate buffered saline (PBS) at 4°C overnight for dissected retinas or eyes. For retinal explants, a shorter fixation time of 3 hr was applied. For all tissues, post fixation we applied 3 x 10 min washes in 1X PBS before the tissues were cryopreserved in 20% sucrose/1X PBS overnight at 4°C. Cryoprotected tissues were embedded in O.C.T™ (Tissue-Tek®, Sakura Finetek U.S.A. Inc., Torrance, CA) and frozen on dry ice. Cryosections were cut at 10 microns on a Leica CM3050s cryostat (Leica Biosystems, Buffalo Grove, IL, USA). Sections were collected on Fisherbrand™ Superfrost™ Plus slides (Thermo Fisher Scientific, Markham, ON).

#### RNAscope and RNA in situ hybridization

RNA in situ hybridization using digoxygenin labeled *Wnt3a* and *Wnt2b* riboprobes was performed as previously described (Alam et al., 2005; Touahri et al., 2015). RNAscope *in situ* hybridization was performed using a RNAscope^®^ Multiplex Fluorescent Detection Kit v2 (ACD; #323110) according to the manufacturer’s directions. ACD probes included: Mm-*Pten* (#316301-C1), Mm-*Dscam* (#834031-C2), Mm-*Th* (#317621-C3), Mm-*Isl1* (#451931-C3) and Mm-*Smpd3* (#815591-C3). Opal^TM^ 520 (Akoya #FP1487001KT; 1:1500), Opal^TM^ 570 (Akoya #FP1488001KT; 1:1500) and OpalTM 690 (Akyoya #FP1497001KT; 1:1500) were used to stain channel 1, 2 and 3 probes. Retinal sections were counterstained with DAPI and mounted in Aquapolymount as described.

#### Immunohistochemistry

Section (Cantrup et al., 2012; Tachibana et al., 2016) and flatmount (Cantrup et al., 2012) immunofluorescence was performed as previously described. The following primary antibodies were used: Acetylated-tubulin (1:500, Abcam #ab11323), ß-catenin (1:500, H-102, Santa Cruz #sc7199), ß-galactosidase (1:400, Chemicon #AB986), Calretinin (1:2000, Swant #76699/4), CD63 (1:200, BD Pharmingen #556019), ChAT (1:500, Chemicon #AB144P), Cholera Toxin β (1:500, Invitrogen/Life #C-34776), Dscam (1:100, R&D Systems #AF3315), Dscam (1:200, R&D Systems #AF3666), Fat3 (1:100, Santa Cruz #sc-133366), Flotillin-1 (1:500, Cell Signaling #3253), γ-tubulin (1:500, Sigma #T3195), GFP (1:500, Abcam #ab5450), Megf10 (1:500, Millipore #ABC10), nSmase2 (1:500, Abcam #ab85017), Pax6 (1/500; Covance Research #PRB-278P), Pten (1:500, Cell Signaling #9559), and TH (1:500, Millipore #AB152). Slides/tissues were washed 3 times in PBS with 0.1% triton X-100 (PBT) and primary antibodies were detected using secondary antibodies conjugated with Alexa568 (1:500, Molecular Probes), Alexa488 (1:500, Molecular Probes), Alexa647 (1:500, Molecular Probes).

#### Electron Microscopy

For electron microscopy (EM) of retinal tissues, eyes were dissected and processed as previously described by us (Cantrup et al., 2012) and others (Zhang et al., 2020). The sections were examined and imaged in a Hitachi H7000 transmission electron microscope at an accelerating voltage of 75 kV. EM with EV samples was performed as followed: purified EVs were fixed on the surface of the carbonated copper grid, followed by counterstaining with uranyl acetate for 1 min. The grids were then briefly rinsed with distilled water, and air dried. The EVs were examined and imaged in a Hitachi H7000 transmission electron microscope at an accelerating voltage of 75 kV.

#### EV purification

For EV purification, eyes were enucleated, the RPE, cornea, lens and blood vessels were removed, and retinas were collected and briefly rinsed with PBS. The retina was placed on a dry 10 cm petri dish, and minced into smaller pieces using a scalpel. The minced tissues were then transferred into a 15 ml corning tube with 10 ml pre-warmed DMEM. The retinal tissues were further dissociated by triturating with a 5 ml pipet. Dissociated retinal cells were incubated in a 37°C water bath for 1 hr with gentle rocking, allowing cells to secrete EVs into the medium. Following the incubation, EVs were isolated using differential centrifugation as previously described (Thery et al., 2006) with the following speeds (300 x *g*, 5 mins; 3024 x *g*, 15 mins; 15,475 x *g*, 15 mins; 153,297 x *g*, 75 mins; 153,297 x *g*, 75 mins). EV pellets were resuspended in 50 μl of cold PBS with 1 x protease inhibitor and 1 x sodium azide and were then further processed for EM imaging or western blotting experiments.

#### Transmission Electron Microscopy

4 µL of purified exosomes were placed on carbon-coated Cu400 TEM grids that were first glow discharged for 30 s (Pelco EasiGlow, Ted Pella Inc.). After 1 min, excess liquid was wicked away, and then washed with 4 µL of distilled water 3 times. 4 µL of 2% uranyl acetate solution was applied for staining for 30 seconds, with excess liquid wicked away before the grids were air-dried. Grids were imaged on a Thermo Fisher Scientific Talos L120C TEM operated at 120 kV using a LaB6 filament in the Microscopy Imaging Laboratory, University of Toronto.

#### Nano-flow cytometry

Amacrine cells were isolated from dissociated P12 retinas using MACS. 1 µl of purified EVs were diluted in 18 µl H_2_O and then labelled with 1 µl anti-CD9 (Santa Cruz; #sc9148) for 30 mins at room temperature. EVs were washed in sterile water, and then stained with Alexa Fluor 647 secondary antibody (Invitrogen, 1:20 dilution) for 20 mins. EVs were washed in sterile water and resuspended in 500 µl sterile water and quantified on the Nanoscale Flow Cytometer (Apogee Flow Systems Inc). EV number is quantified from scatterplots of Alexa 647 emission (CD9^+^ levels) and long angle light scatter (LALS), for size distribution). EVs were defined particles greater than 100 nm in diameter.

#### Western blotting and silver staining

Retinas were lysed in in NP-40 lysis buffer (0.05 M Tris, pH 7.5, 0.15 M NaCl, 1% NP-40, 0.001 M EDTA) with protease (1X protease inhibitor complete, 1 mM PMSF), proteasome (MG132 at 0.05 mM) and phosphatase (50 mM NaF, 1 mM NaOV) inhibitors. 10 μg of lysate from whole cells or tissue was run on SDS-PAGE gels for western blot analysis as described previously (Ma et al., 2007). To characterize purified EVs, they were boiled at 95°C for 2 mins to break the EV membrane. The protein concentrations were determined using a Micro BCA™ Protein Assay Kit (ThermoFisher Scientific, #23235) following the manufacturer’s instructions. 2 μg of lysate was incubated with the blots overnight at 4°C. Primary antibodies included: β-actin (1:10000, Abcam #8227), Flotillin-1 (1:1000, Cell Signaling #3253), Megf10 (1:1000, Millipore #ABC10), Pten (1:1000, Cell Signaling #9559), ß-Catenin-non-phospho (Active) (Ser33/37/Thr41) (D13A1) (1:1000; Cell Signaling #8814) and TSG101 (1:1000, Sigma #HPA006161). Blots were washed 3 x 10 min in TBST before incubating in secondary antibodies, including either Goat Anti-Rabbit IgG (H+L)-HRP Conjugate (1/50,000; Bio-Rad #1721019) or Goat Anti-Mouse IgG (H+L)-HRP Conjugate (1/50,000; Bio-Rad #1721011). Western blot signals were converted to a chemiluminescent signal using an Amersham ECL Prime Western Blotting Detection Reagent (Cytiva #RPN2232) according to the manufacturer’s instructions. Signal was detected by exposing to autoradiography film (LabForce #1141J52) or using a Bio-rad gel doc and GelCapture MicroChemi 2.2.0.0 software. Each western blot was performed on lysates from three independent retinas, and densitometries were calculated using UN-SCAN-IT gel densitometry software (Silk Scientific). Absolute values (“relative”) or normalized values (over ß-actin) were plotted as indicated.

#### Mass Spectrometry

EVs were lysed in NP-40 lysis buffer and 2 μg of total protein was separated on a 10% SDS-PAGE gel and silver stained. Proteins were digested using trypsin as described (Chaturvedi et al., 2012). Peptide extracts were concentrated by Vacufuge (Eppendorf). Peptides were analyzed by LC-MS/MS using a Dionex Ultimate 3000 RLSC nano HPLC (Thermo Scientific) and Orbitrap Fusion Lumos mass spectrometer (Thermo Scientific) at the Ottawa Hospital Research Institute Proteomics Core Facility (Ottawa, Canada) as described (Chaturvedi et al., 2012). MASCOT software version 2.6 (Matrix Science, UK) was used to infer peptide and protein identities from the mass spectra. The observed spectra were matched against mouse sequences from SwissProt (version 2016-09) and also against an in-house database of common contaminants. Results were then exported to Scaffold version 4.8.2 (Proteome Software, USA) for further analysis. Briefly, proteins identified with less than 2 peptides and proteins with a probability of identification of less than 95% were filtered out. REACTOME enriched pathway clustering was obtained using pathfind R Bioconductor package (Ulgen et al., 2019), extracting four types of plots: 4 types of plots: 1) Simple dot plot for top 15 pathways; 2) Clustered dot plot that shows that clusters together similar pathways; 3) Term-gene enrichment map; 4) UpSet plot that shows genes in enriched pathways. For gene ontology analysis, we present Scatter plots that reflect semantic similarities and treemaps using a rrvgo Bioconductor package https://bioconductor.org/packages/release/bioc/vignettes/rrvgo/inst/doc/rrvgo.html.

#### scRNA-seq analysis

10x scRNA-seq data of mouse developing retina was obtained from GSE118614 (Clark et al., 2019). Further processing and analyses were performed with the Seurat v.3.2.3 R package (Butler et al., 2018). Low quality cells were excluded by filtering out cells with fewer than 500 detected genes and cells with mitochondrial RNA more than 5%. The data was then transformed by the SCTransform function while regressing out the variance due to mitochondrial RNAs. Clustering was performed by the RunPCA, FindNeighbors and FindClusters functions using the first 30 principal components. The 2D projection of the clustering was carried out by the RunUMAP function. The annotation of cell type to each cluster was performed by using the same set of markers as in (Clark et al., 2019). Expression of selected genes was plotted using the FeaturePlot function.

#### Retinal explants

Eyes were removed from P0 CD1 pups using surgical forceps and placed in 1x PBS, pH 7.4 on ice. The eye was then held by the cornea to remove the iris, lens and anterior segment. The retina was then removed from the underlying retinal pigment epithelium (RPE), choroid and sclera. Isolated retinas were transferred to a sterile 10 cm dish in cold 1x PBS pH 7.4 for further processing. Retinas were flatmounted onto 13 mm, 0.8 μm Nuclepore Track-Etch membrane (GE Healthcare, catalog number: 110409) and floated on 2 ml of retinal explant media (50% DMEM, 25% HBSS, 25% heat-inactivated horse serum, 200 μM L-glutamine, 0.6 mM HEPES, 1% Pen-Strep) in individual wells of a 24-well plate. Recombinant Wnt3a protein (Catalog # 5036, R&D Systems, final concentration of 100 ng/ml in water), XAV939, a Wnt antagonist (Catalog # S1180, Selleckchem; final concentration of 10 mM in DMSO), or DMSO was added to the culture media. Half the media was removed and replaced with fresh media containing drugs every second day.

#### Cell spacing analysis

Flat-mounted retinas or retinal explants were imaged through the z axis to ensure all ChAT^+^ and Islet^+^ cells were included in the images. ImageJ was used for collecting the X-Y coordinates of ChAT^+^ and Islet1^+^ amacrine cells. These values were imported into a customized program that computes the Voronoi tessellation of the field and the nearest neighbour distances of each individual cell, from which the regularity indices of the distribution of Voronoi areas and nearest neighbour distances were calculated (mean/standard deviation). Explant cultures were also assessed for minimal spacing between cells within a field, expressed as the effective radius (ER) derived from the density recovery profile (Rodieck, 1991). Border cells with uncertain spatial statistics were excluded from the measurements, all as previously described (Keeley et al., 2020; Raven and Reese, 2002).

#### Imaging

Digital images were captured using fluorescent light from a Leica DM18 inverted microscope and Leica Application Suite X (LASX) software for cell counts. Confocal images were captured using a Nikon A1 laser scanning confocal microscope available at the Centre for Cytometry and Scanning Microscopy at Sunnybrook. Adobe Photoshop 2021 was used to make Figures. A license to BioRender.com or the software Inkscape 0.48 was used to prepare schematics.

#### Statistical analysis

Statistical analysis was performed on a minimum of three animals per each genotype (N; biological replicates) with a minimum of three sections from each animal (n; technical replicates). Individual dots in the plots indicate a single biological replicate (N) after averaging the three technical replicates (n). All statistical analysis and graph designs were performed using GraphPad Prism Software version 8.0 (GraphPad Software). Error bars represent standard error of the mean (SEM). Unpaired two tailed Student’s t test was used to calculate statistical significance between two experimental groups. For multiple comparisons of more than two groups, a one-way ANOVA was used followed by a Tukey post hoc-test. For the analysis of Dscam puncta distribution, a G-test was applied. In all plots, statistical significance was defined as a p-value less than 0.05, with p values denoted as follows: p<0.05 *, <0.01 **, <0.001 ***.

## SUPPLEMENTAL INFORMATION

**Supplementary Figure 1.**
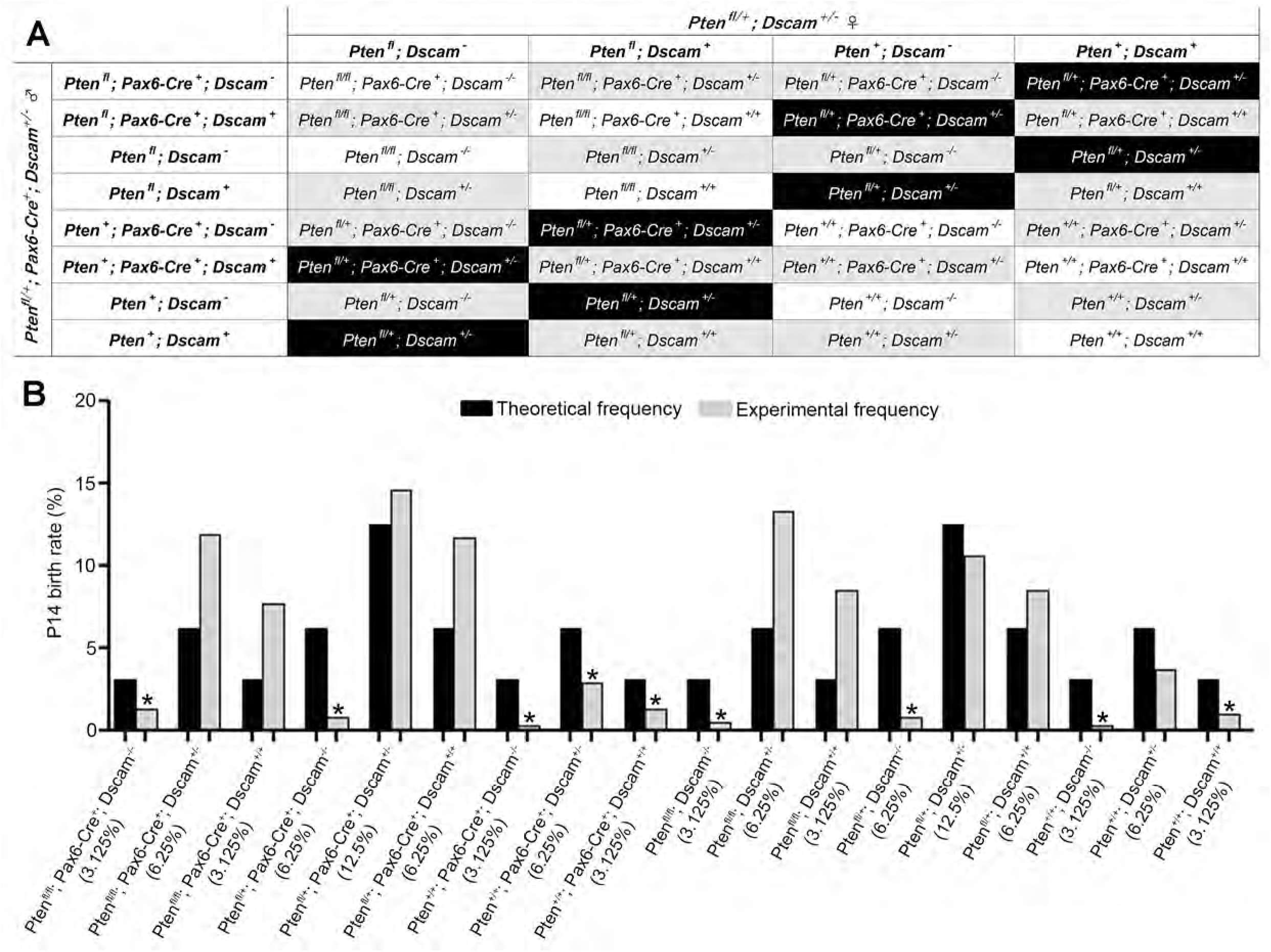
*Dscam*^del17^ and *Pten*^cKO^*;Dscam*^del17^ pups are underrepresented in double heterozygous intercrosses. (A) Punnett square showing all possible genotypes of the progeny from *Pten^fl/+^; Dscam^+/del17^* female and *Pten^fl/+^; Pax::Cre^+^;Dscam^+/del17^* male intercrosses. 56 litters were analysed for a total of 376 live P14 pups. White, grey and black boxes indicate the genotypes with birth rate of 3.125%, 6.25%, and 12.5%, respectively. (B) Graph comparing the theoretical birth rates of each genotype with the experimental birth rates. Black bars represent the theoretical frequency, and grey bars represent the experimental birthrates. Asterisk indicates the genotypes with lower experimental birth rate compared to theoretical birth rate.

**Supplementary Figure 2.**
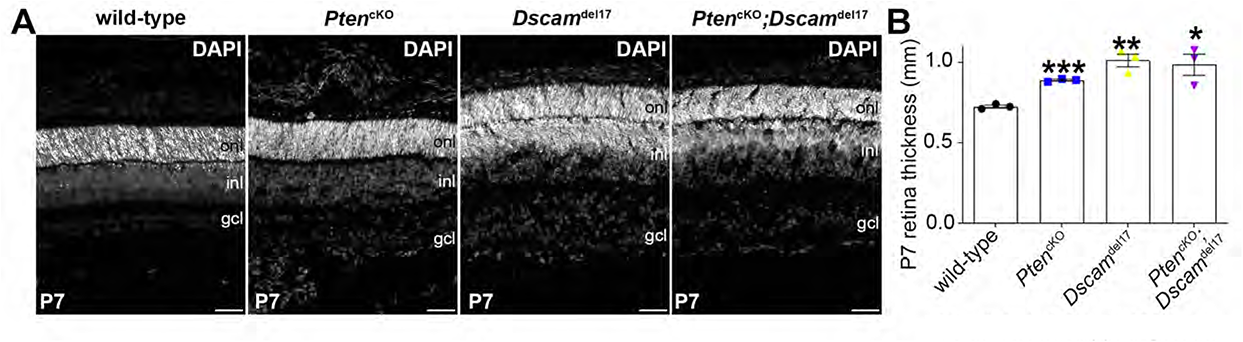
Increased retinal thickness in *Pten*^cKO^ and *Dscam*^del17^ single and double mutant retinas. (A) DAPI staining of transverse sections through the retinas of P7 wild-type, *Pten* ^cKO^, *Dscam*^del17^, *Pten*^cKO^*;Dscam*^del17^(D) retinas. (B) Measured thickness of retinas in each group. p values are denoted as follow: <0.05 *, <0.01 **, <0.001 ***. Scale Bar: 200μm.

**Supplementary Figure 3.**
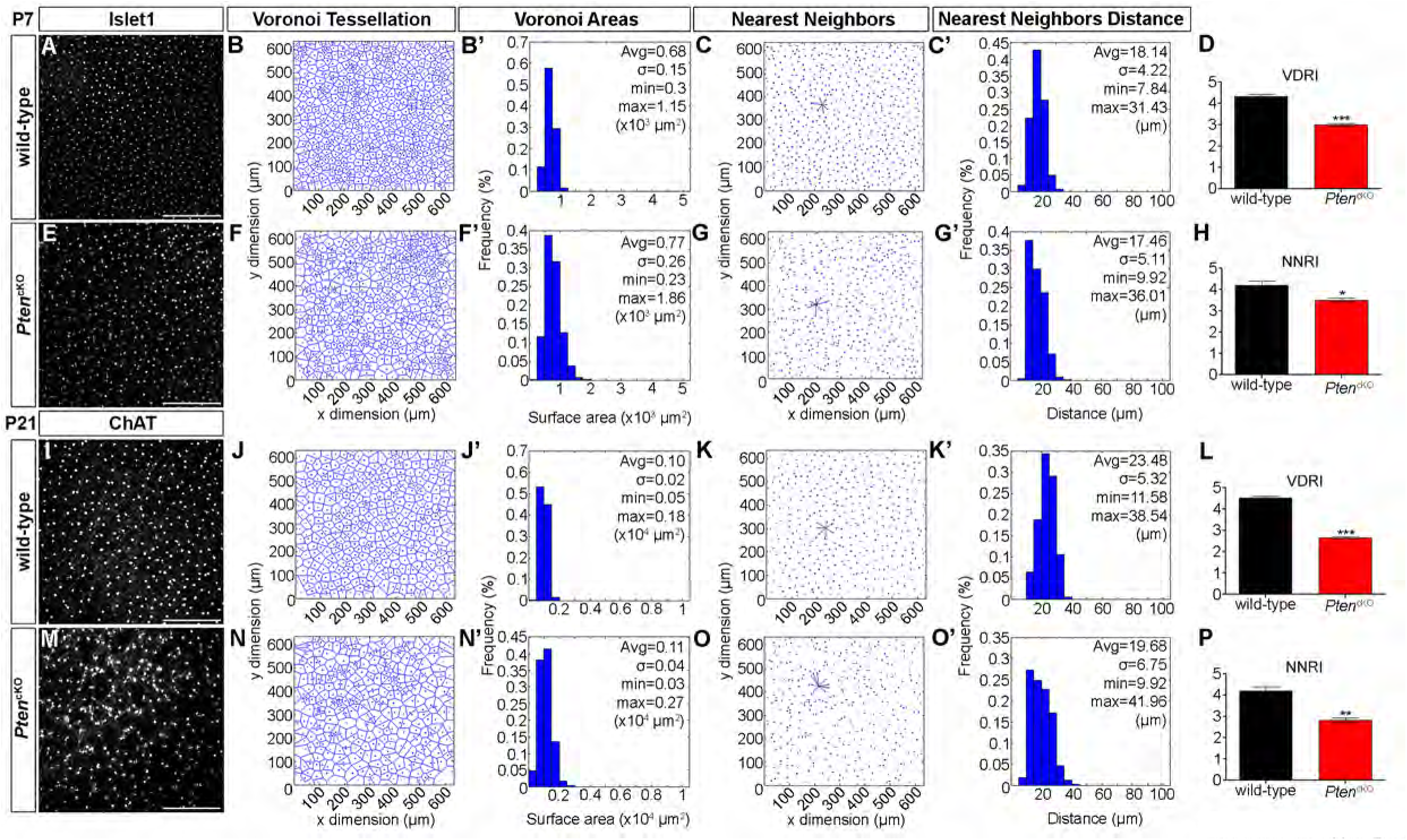
Irregular cholinergic mosaics in *Pten*^cKO^ retinas at P7 and P21. (A-H) Immunolabeling of P7 wild-type (A), and *Pten*^cKO^ (E) retinal flatmounts for Islet1. Voronoi tessellation and Voronoi areas depicting the distribution of Islet1^+^ amacrine cells in P7 wild-type (B,B’) and *Pten*^cKO^ (F,F’) retinas. Nearest neighbors and frequency distributions of nearest neighbor distances of Islet1^+^ reference cells in P7 wild-type (C,C’) and *Pten*^cKO^ (G,G’) retinas. Calculation of Voronoi domain (D) and Nearest Neighbor (H) regularity indices for Islet1^+^ amacrine cells in the wild-type and *Pten*^cKO^ retinas. (I-P) Immunolabeling of P21 wild-type (I), and *Pten*^cKO^ (M) retinal flatmounts for ChAT. Voronoi tessellation of ChAT^+^ amacrine cells in P21 wild-type (J,J’) and *Pten*^cKO^ (N,N’) retinas. Nearest neighbors and frequency distributions of nearest neighbor distances of ChAT^+^ reference cells in P21 wild-type (K,K’) and *Pten*^cKO^ (O,O’) retinas. Calculation of Voronoi domain (L) and Nearest Neighbor (P) regularity indices for ChAT^+^ amacrine cells in P21 wild-type and *Pten*^cKO^ retinas. p values are denoted as follow: <0.05 *, <0.01 **, <0.005 ***. Scale Bar: 200μm.

**Supplementary Figure 4.**
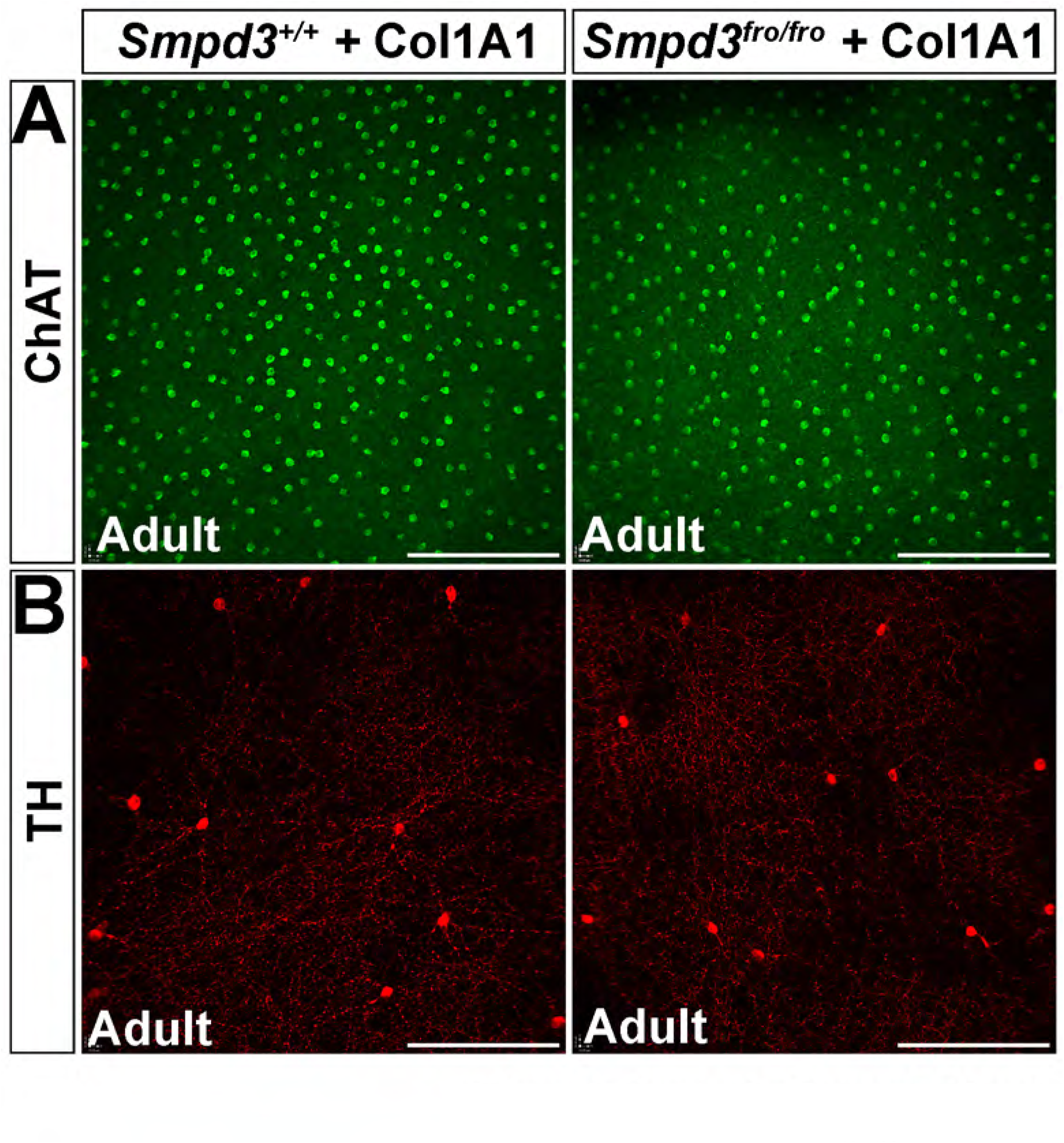
*Fro/fro (Smpd3)* null mutant mice have normal dopaminergic and cholinergic amacrine cell mosaics. (A,B) Immunolabeling of adult wild-type and *Smpd3^fro/fro^*+Col1A1 flatmount retinas with ChAT (A) and TH (C). Scale Bar: 200μm.

### Supplementary Tables

**Supplementary Table 1. ClueGO analysis of proteins enriched in P14 wild-type and *Pten*^cKO^ retinal EVs.** Duplicates were removed.

**Supplementary Table 2. Biological processes enriched in P14 wild-type retinas.**

**Supplementary Table 3. Biological processes enriched in P14 *Pten*^cKO^ retinas.**

**Supplementary Table 4. Reactome pathways enriched in P14 wild-type retinas.**

**Supplementary Table 5. Reactome pathways enriched in P14 *Pten*^cKO^ retinas.**

